# RNA Pol II pausing facilitates phased pluripotency transitions by buffering transcription

**DOI:** 10.1101/2022.04.21.489065

**Authors:** Abderhman Abuhashem, Alexandra G. Chivu, Yixin Zhao, Edward J. Rice, Adam Siepel, Charles G. Danko, Anna-Katerina Hadjantonakis

## Abstract

Promoter-proximal RNA Pol II pausing is a critical step in transcriptional control. Pol II pausing has been studied predominantly in tissue culture systems. While Pol II pausing has been shown to be required for mammalian development, the phenotypic and mechanistic details of this requirement are unknown. Here, we find that loss of RNA Pol II pausing stalls pluripotent state transitions in the epiblast of the early mouse embryo. Using *Nelfb^-/-^* mice and a novel NELFB- degron mouse embryonic stem cells, we show that mouse ES cells (mESCs) representing the naive state of pluripotency successfully initiate a transition program, but fail to balance levels of induced and repressed genes and enhancers in the absence of NELF. Consistently, we find an increase in chromatin-associated NELF during pluripotency transitions. Overall, our work reveals the molecular and phenotypic roles of Pol II pausing in pluripotency and introduces Pol II pausing as a modulator of cell state transitions.

## INTRODUCTION

Transcriptional regulation is a hallmark of cell fate specification (Cramer 2019; Johnston and Desplan 2010). Upstream cell extrinsic inputs, such as growth factor signaling, mediate cell intrinsic responses which converge on the transcriptional machinery to regulate recruitment of RNA polymerase II (Pol II) at specific gene targets, and thereby, gene expression (Adelman and Lis 2012; Core and Adelman 2019; Pope and Medzhitov 2018). Pol II promoter-proximal pausing (Pol II pausing) has been identified as a key rate-limiting step of transcription in metazoans (Core and Adelman 2019; Shao and Zeitlinger 2017). Pol II pausing represents a brief halt of transcription 30-60 nucleotides downstream of the transcription start site (TSS). This pause is regulated by two protein complexes, the DRB-sensitivity inducing factor (DSIF) and the Negative Elongation Factor (NELF) (Chen et al. 2018; Yamaguchi et al. 1999). Releasing paused Pol II is achieved by phosphorylation of NELF, DSIF, and Pol II by CDK9 (Adelman and Lis 2012). These phosphorylation events result in the dissociation of NELF and progression of DSIF and Pol II into productive elongation.

The functional role of Pol II pausing has been studied in a variety of contexts, predominantly *in vitro*. Genomic and structural studies have revealed that the paused Pol II sterically hinders new initiation events, and NELF occupies a large interaction surface with Pol II which is substituted for elongation factors, such as the PAF complex, upon pause-release (Gressel et al. 2019; Shao and Zeitlinger 2017; Vos et al. 2018b, 2018a). Kinetically, the stability of the paused polymerase, estimated at a time scale of minutes, highlights the importance of regulating this step (Gressel et al. 2019; Krebs et al. 2017; Shao and Zeitlinger 2017; Steurer et al. 2018). Several transcription factors and signaling components can act specifically on the pause-release step to regulate gene expression (Danko et al. 2013; Gilchrist et al. 2012; Liu et al. 2015; Williams et al. 2015; Yu et al. 2020; Henriques et al. 2013). These include the heat shock response, and glucocorticoid, TGF-ß and ERK singling pathways. Attempts to perturb pausing have been achieved primarily via loss of function studies of NELF proteins, which play an exclusive role in Pol II pausing but not elongation (Chen et al. 2018). These studies have revealed that NELF is required for early development in *Drosophila*, Zebrafish, and mice (Amleh et al. 2009; Wang et al. 2010; Yang et al. 2016; Abuhashem et al. 2022). Despite several studies revealing broad requirements of NELF in development and a variety of tissue-specific contexts in mice, the underlying molecular mechanisms have remained largely unknown.

Development represents a dynamic period of gene regulation where cells must constantly change their gene expression patterns as they adopt new states (Johnston and Desplan 2010). Consistent with this notion, NELF knockout mice show embryonic lethality at peri-implantation stages (Amleh et al. 2009; Williams et al. 2015). Given the advantage of mouse embryonic stem cells (mESCs), the in vitro counterpart of the pluripotent epiblast, to model key aspects of early mouse development, NELF knockout and knockdown studies in mESCs revealed that Pol II pausing is essential for cellular differentiation (Amleh et al. 2009; Williams et al. 2015). However, interpretation of these results has been complicated due to potential secondary effects resulting from long term NELF knockout and compounding proliferation defects (Williams et al. 2015). Additionally, the cellular and molecular details of the developmental arrest of embryos remain unclear.

In this study, we perform a comprehensive characterization of the role of NELF in early mouse development, with a focus on pluripotent cell state transitions. We utilized a *Nelfb* knockout mouse model to show that *Nelfb^-/-^* embryos exhibit normal pre-implantation development as they were recovered at Mendelian ratios with cell lineage specification comparable to wild type embryos. We show that pre-gastrulation lineages are properly assigned, except for the posterior epiblast, and that mutant embryos fail pre-gastrulation ∼E5.75. The epiblast lineage is specified during the blastocyst stage, at ∼E3.5, and transitions from a naïve state in the blastocyst to a primed state prior to gastrulation at ∼E6.5 in a sequential manner (Morgani et al. 2017). To further investigate the molecular basis of the defect observed in embryos, we took advantage of mESCs as a paradigm that models pluripotency transitions from the naïve to the formative and primed states (Hayashi et al. 2011). To allow efficient, rapid and reversible depletion of NELFB protein, we designed a homozygous knock-in *Nelfb* degron allele using the dTAG system (Nabet et al. 2018). This model recapitulated the defects of pluripotency transitions and priming observed in *Nelfb^-/-^* embryos and highlighted a requirement of NELFB during pluripotency transitions in mESCs.

To gain further mechanistic insights into the defect observed within the epiblast layer of the embryo, we used the mESC model and coupled chromatin immunoprecipitation followed by sequencing (ChIP-seq) and nascent transcriptomic analyses (PRO-seq) which showed widespread binding of NELF at both promoters and enhancers, in support of previous reports (Core et al. 2012; Henriques et al. 2018). Our NELFB degron cells enabled acute degradation of NELF in mESCs which resulted in global loss of Pol II pausing at both gene promoters as well as enhancers within 30 minutes. Surprisingly, degrading NELF transiently in the context of pluripotency transitions from naïve to the formative state caused a hyper-induction of genes associated with the formative state accompanied by hyper-silencing of downregulated genes. This is in agreement with recent studies suggesting that absence of NELF perturbs fate transition events more so than steady-state cellular function in a variety of contexts (Yu et al. 2020; Robinson et al. 2021; Hewitt et al. 2019). Accordingly, we observed increased recruitment of NELF to chromatin during pluripotency state transitions. Our data leads us to propose a model whereby Pol II pausing facilitates state transitions by attenuating and buffering the expression of genes associated with cell identity, thereby enabling coordinated transitioning between cell states.

## RESULTS

### *Nelfb^-/-^* embryos display defects in pluripotent epiblast state transitions

*Nelfb^-/-^* mouse embryos exhibit embryonic lethality at post-implantation stages (Amleh et al. 2009). To further characterize the defects observed in *Nelfb^-/-^* embryos, we utilized a mouse model that harbors a deletion of the first four exons of *Nelfb* resulting in a protein-null allele (Figure S1A)(Williams et al. 2015). Since previous reports suggested that *Nelfb^-/-^* blastocyst- stage embryos might show defects in cell fate specification, we initiated our analysis at pre-implantation stages of embryonic development (Amleh et al. 2009; Williams et al. 2015). We collected early (E3.25) to late (E4.5) stage blastocysts and immunostained for lineage specific markers: NANOG, GATA6, and CDX2 to identify the pluripotent epiblast (Epi), primitive endoderm (PrE), and trophectoderm (TE) lineages, respectively. *Nelfb^-/-^* blastocysts were morphologically indistinguishable from heterozygous littermates and displayed the correct spatial distribution of its three cell lineages (Figure 1A). To assess the developmental progression of blastocysts, we staged them based on total cell number per embryo as an accurate metric of stage, and assigned a lineage identity to each cell based on its relative expression of markers (Lou et al. 2014; Saiz et al. 2016b, 2016a). *Nelfb^-/-^* embryos did not exhibit a defect in total cell number, ratio of TE, Epi or PrE, or the gradual assignment of the inner cell mass cells (NANOG/GATA6 double-positive, or DP) to epiblast and primitive endoderm fates (Figure 1B, S1B and S1C). Moreover, we found that *Nelfb^-/-^* embryos could be recovered at Mendelian ratios up until post-implantation, but show significant defects by E7.5 (Figure 1C and S1D)(Amleh et al. 2009). Thus, our analysis of pre-implantation stage mutant mouse embryos suggests the *Nelfb* is dispensable for cell lineage specification, survival, and implantation of blastocysts.

**Figure 1.**
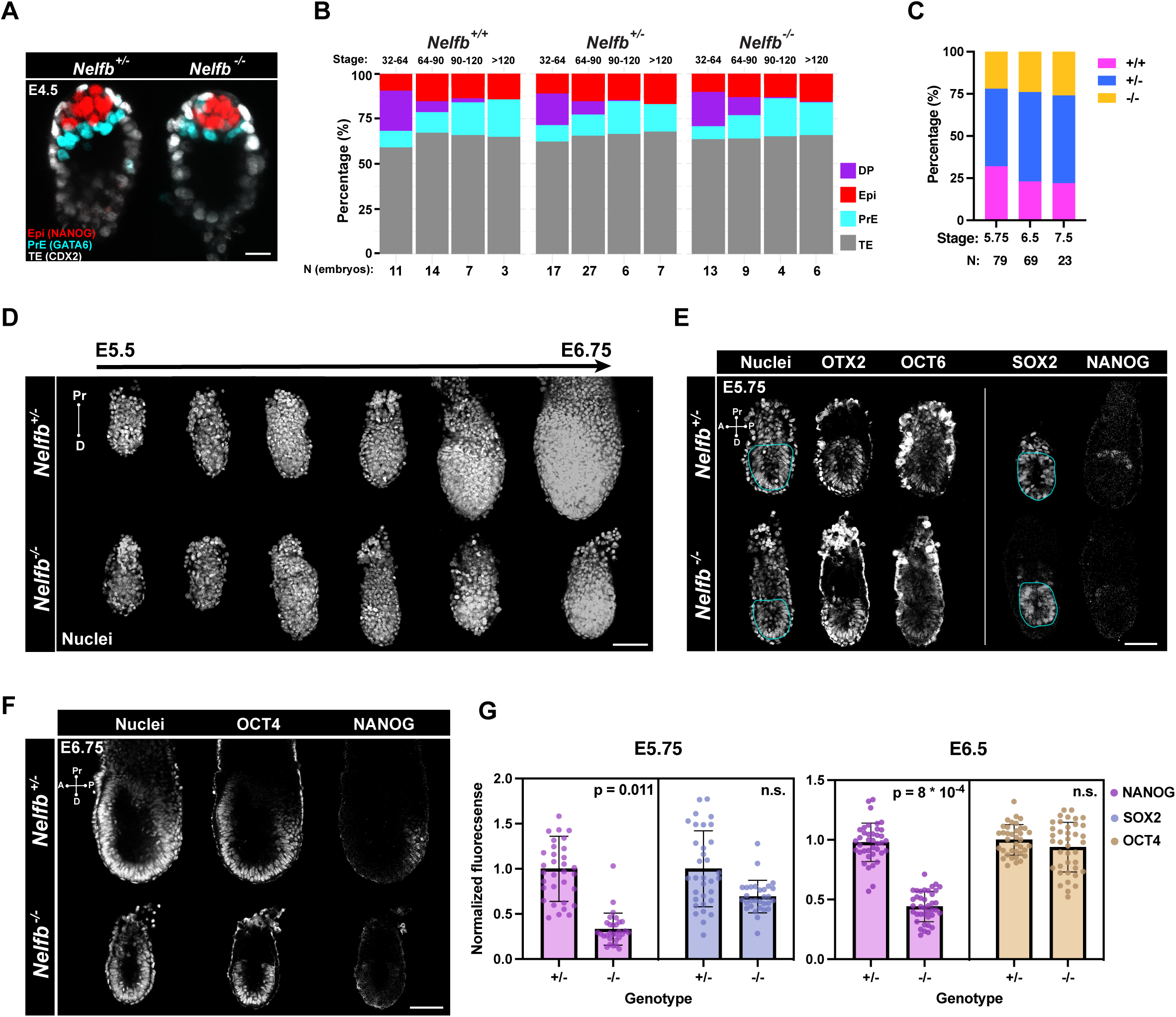
*Nelfb^-/-^* embryos display defects in pluripotent epiblast state transitions. (A) Immunofluorescence of E4.5 blastocysts labeling epiblast: NANOG, primitive endoderm: GATA6, and trophectoderm: CDX2. Several Z slices are shown in maximum intensity projection (MIP) to show the ICM. Scale bar 15μm. (B) Stacked bar plot representing percentage of each lineage in blastocysts sorted by stage: total cell number per blastocyst, and genotype. (C) Stacked bar plot representing percentage of each *Nelfb* genotype at different post-implantation stages. (D) Maximum Intensity Projection (MIP) of embryos dissected at stages between E5.5 and E6.75 at 0.25 increments. Nuclei are shown to reflect whole embryo. Nuclei were labeled with Hoechst. Scale bar 100 μm. (E) Immunofluorescence of E5.75 embryos of select pluripotency markers. The bordered region highlights the epiblast cup. The vertical line means separate embryos. Single Z slices are shown. Scale bar 50 μm. (F) Immunofluorescence of E6.75 embryos of select pluripotency markers. Nuclei were labeled with Hoechst. Single Z slices are shown. Scale bar 100 μm. (G) Normalized immunofluorescence intensity per epiblast nuclei for pluripotency markers. Single dots are single nuclei. Quantifications show four embryos per group. Statistical testing using t-test was performed on embryo averages. Error bars show standard deviation. p < 0.05 was used to determine significance.

To determine when development became dysregulated in *Nelfb^-/-^* mutants, we collected post-implantation stage embryos prior to and after the onset of gastrulation (E5.5-E6.75). By E6.75 *Nelfb^-/-^* embryos were smaller than their wild-type or heterozygous littermates (Figure 1D). Prior to this stage at E5.75, *Nelfb^-/-^* embryos did not display proliferation or size defects, assayed by staining for phosphorylated H3 and measuring the epiblast section area, respectively (Figure S1E and S1F). To determine whether cell fate specification was affected, we analyzed the expression and distribution of lineage specific transcription factors for epiblast (SOX2), visceral endoderm (GATA6), and extraembryonic ectoderm (CDX2). All three lineages were present, with cells organized in the expected spatial arrangement (Figure S1G and S1H). We next crossed *Nelfb^+/-^* mice to the *Afp-GFP^Tg^* visceral endoderm and *Hex-tdTomato^Tg^* anterior visceral endoderm reporters (Kwon et al. 2006; Wu et al. 2017). Visualization of these lineage specific reporters revealed that *Nelfb^-/-^* embryos possessed a visceral endoderm layer and had successfully specified the distal/anterior visceral endoderm population that was able to migrate anteriorly (Figure S1H and S1I). These results suggest that at E5.75, when the anterior visceral endoderm has completed its migration and prior to the onset of gastrulation, *Nelfb^-/-^* embryos are indistinguishable from their wild-type and heterozygous littermates, by morphology, and lineage specific marker expression and localization.

Given previous reports suggesting the *Nelfb^-/-^* mESCs, the *in vitro* counterpart of the epiblast of the embryo, show defects in differentiation, we went on to examine the epiblast population further. Pluripotent epiblast cells are specified in the mid-to-late blastocyst, and subsequently progress through pluripotent state transitions before they exit pluripotency and differentiate at gastrulation (Morgani et al. 2017). Distinct stages in the pluripotency continuum include the early naïve state (E4.5, NANOG+), the subsequent formative state (E5.5, NANOG-, OTX2+), and posterior primed state (posterior epiblast at E5.75-E6.5, NANOG+ OTX2+). At E5.75, we found that cells of the epiblast of *Nelfb^-/-^* embryos successfully induced expression of the formative state markers OTX2 and OCT6 (Figure 1E). However, mutant embryos lacked a weak NANOG+ population representing the posterior primed state. By E6.75, the posterior primed population expressed NANOG robustly and surrounded the primitive streak in heterozygous and wild-type embryos but remained largely absent in *Nelfb^-/-^* despite expression of comparable levels of the pan pluripotency marker OCT4 (Figure 1F and 1G). Subsequently, mutant embryos fail to induce a T+ primitive streak marking gastrulation at E6.75 (Figure S1J). These results show that *Nelfb^-/-^* embryos exhibit defects at early post-implantation stages (after the AVE has migrated but before the onset of gastrulation, E5.75), where cells of the posterior epiblast are unable to attain a posterior primed state and progress to gastrulation.

### NELFB-depleted mESCs recapitulate defects in pluripotent state transitions observed in the embryo

To characterize the pluripotency transition defects observed in mutant mouse embryos at the molecular level, we sought to develop an *in vitro* model of NELFB loss in mESCs. mESCs can be cultured under defined conditions in the presence of FGF and ACTIVIN to model pluripotency transitions to the subsequent formative and primed states (Hayashi et al. 2011; Morgani et al. 2017). We failed to derive mESCs from *Nelfb^-/-^* embryos, consistent previous reports (Williams et al. 2015). Although previous studies of NELFB in cell culture models used either knockdown or conditional knockout systems, these methods require days to achieve successful depletion or deletion, resulting in an inability to discern primary versus secondary effects (Wu et al. 2020). We therefore took advantage of the recently developed dTAG protein degron system (Nabet et al. 2018). By fusing a protein of interest to a tag, FKBP12^F36V^, the target protein can be acutely and reversibly degraded using a heterobifunctional small molecule, such as dTAG-13, that targets FKBP12^F36V^ for proteasomal degradation (Figure 2A).

**Figure 2.**
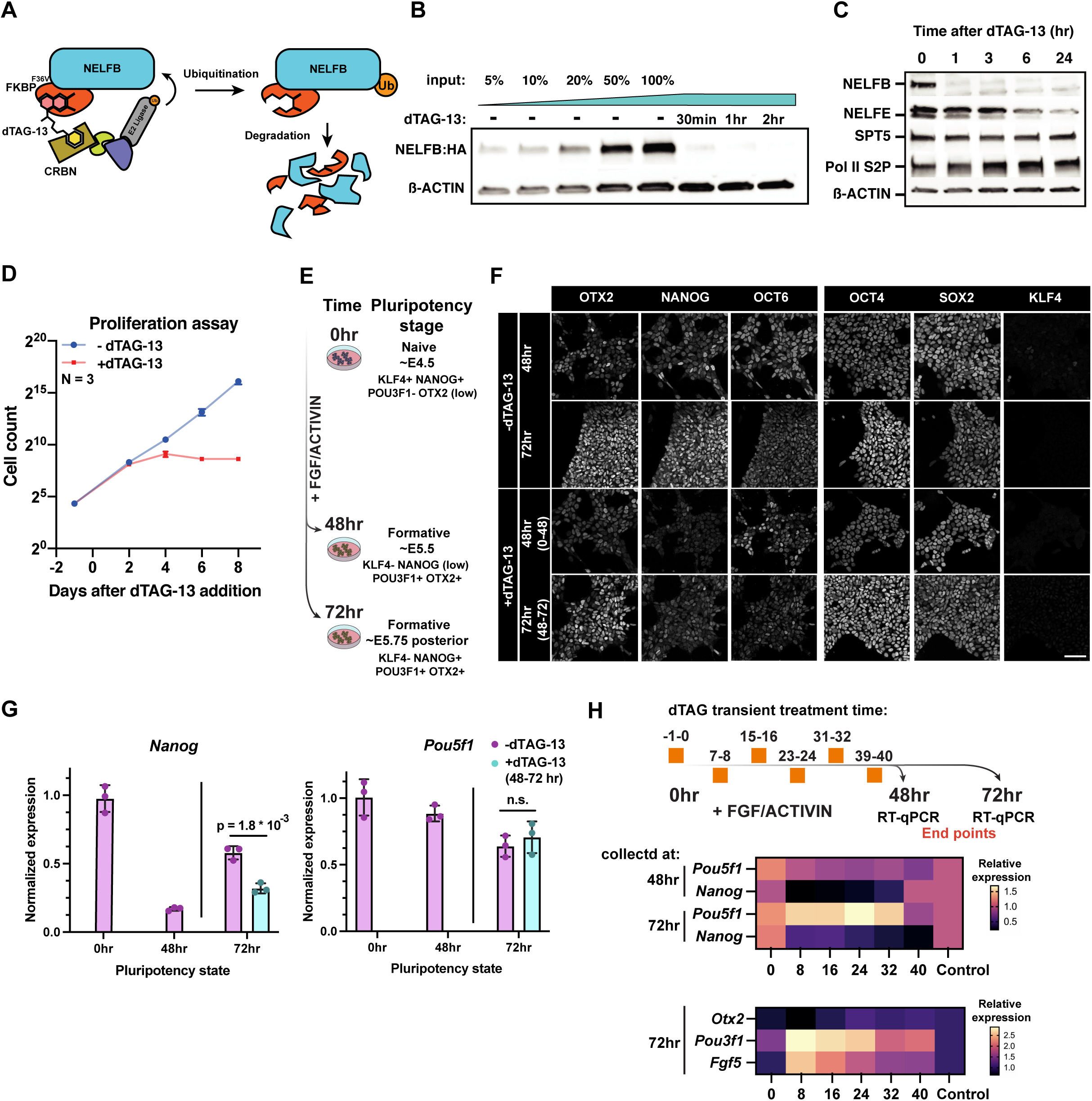
NELFB-depleted mESCs recapitulate defects in pluripotent state transitions observed in the embryo. (A) Schematic of the dTAG targeted protein degradation system. (B) Western blot of NELFB degradation efficiency and dynamics following 500nM dTAG-13 treatment. Input refers to relative amount of protein loaded to the gel. (C) Western blot of transcription associated proteins following NELFB degradation for varying time periods. (D) Proliferation assay of *Nelfb^deg^* mESCs in the presence and absence of 500nM dTAG-13. Cells were counted and passaged every two days. (E) Schematic of the pluripotency transitions protocol *in vitro*. The schematic shows corresponding *in vivo* stages and markers expression. (F) Immunofluorescence of *Nelfb^deg^* mESCs following pluripotency transitions with and without dTAG-13 at 48 and 72 hrs. The time interval in parentheses in the treatment panels refers to the time of adding dTAG-13. Scale bar 50 μm. (G) Normalized RT-qPCR of select factors from experiment in (F). The +dTAG-13 marks the addition of dTAG-13 between hours 48-72 of pluripotency transitions. Data was normalized to *Actb* levels. Statistical testing using t-test was performed on embryo averages. Error bars show standard deviation. p < 0.05 was used to determine significance. (H) (top) Schematic of experiment showing different times of adding dTAG-13 for 1 hour followed by washing. Each timepoint represents one condition. Cells were collected for RT-qPCR at hour 48 and 72 of transitions. (middle) Heatmap of normalized RT-qPCR expression relative to control. Naïve factors are shown. (bottom) Heatmap of normalized RT-qPCR expression relative to control. Formative factors are shown.

We generated a *Nelfb-FKBP12^F36V^-2xHA* homozygous knock-in mESC line (the homozygous line hereafter is referred to as *Nelfb^deg^*) using CRISPR editing with homology directed repair (HDR) (Figure S2A, S2B and S2E)(Ran et al. 2013). We noted that our system is capable of degrading NELFB to undetectable levels within 30 mins of adding the degradation inducing small molecule dTAG-13 (Figure 2B and S2C). Upon dTAG-13 washing, NELFB levels recovered significantly within 3-5 hours (Figure S2D). Notably, NELFB degradation did not affect the levels of other transcription machinery proteins such as SPT5 and Pol II S2P (Figure 2C). However, NELFE levels were markedly reduced 24 hours after inducing degradation as expected given the interdependence between the NELF complex proteins (Figure 2C)(Narita et al. 2007). The cells did not display any toxicity to the edited allele or to dTAG-13 treatment in the absence of the edited allele, as assayed by their proliferation capacity (Figure S2F). Continuous degradation of NELFB resulted in reduced proliferation following 3-4 days, and did not affect the expression of pluripotency markers, as studies reported (Figure 2D and S2G)(Williams et al. 2015; Amleh et al. 2009). These data demonstrate that the NELFB degron system in mESCs achieves specific, rapidly inducible, and reversible protein depletion.

To assess whether NELFB depletion can affect transitions between pluripotent states *in vitro*, we utilized a protocol for directing mESCs representing the naive state of pluripotency into EpiLC representing a subsequent formative/primed pluripotent state (Hayashi et al., 2011). Naïve mESCs were maintained in naïve conditions – N2B27 + 2i (MEK and GSK-3β inhibitors) + LIF – and transferred to N2B27 + FGF2 + Activin for 48 to 72 hours to induce pluripotency transitions (Figure 2E). By 48 hours, cells had downregulated KLF4 and NANOG, markers associated with the naïve state of pluripotency, and activated expression of formative pluripotency markers, OTX2 and OCT6 (Figure 2F). At 72 hours, cells maintained formative markers, while upregulating NANOG, consistent with a posterior-like primed pluripotent state (Figure 2F). Degron-induced NELFB depletion from 0-72 hour did not affect the onset of formative markers expression but resulted in a marked loss of NANOG at 48 hours without subsequent upregulation at 72 hours (Figure 2F and S2H). Given that continuous degradation of NELFB from 0-72 hours resulted in reduced proliferation, we degraded NELFB for a 24-hour window, at 48-72 hours of FGF + ACTIVIN exposure (posterior priming phase). Under these conditions, we found that cells recapitulated the failure in reactivation of NANOG, without affecting cell proliferation at 72 hours (Figure 2F and 2G). These data demonstrate that we generated a system that faithfully recapitulates our *in vivo* findings in embryos *in vitro* in a mESCs model, with a fine temporal control that can uncouple acute from secondary effects of NELFB loss. Furthermore, these data define a 24-hour time window when NELFB is required within the epiblast and reveal that acute loss of NELFB specifically affects epiblast cells as they transition between OTX2+ OCT6+ NANOG- and subsequent OTX2+ OCT6+ NANOG+ states.

To further define the time requirement of NELFB during this process, we took advantage of the reversibility of our degradation system. mESCs were cultured in the presence of FGF + ACTIVIN to transition them from naïve to formative pluripotent stages with one hour degradation followed by washing at 0, 8, 16, 24, 32, 40 hours (for example, treating with dTAG-13 between -1 and 0 hours, or 7 and 8 hours, and so on). Samples were collected for RT-qPCR analyses at 48 and 72 hours. We found that treatments at 16, 24, 32 hours had the strongest effect on *Nanog* expression at both 48 and 72 hour time points, with no change when degradation occurred at 0 hour immediately before starting the transitions (Figure 2H and S2I). Concomitantly, certain formative markers, including *Fgf5* and *Pou3f1*(encoding OCT6 protein), were further upregulated at the same timepoints, with little to no change to the expression level of the pan-pluripotency marker *Pou5f1* (encoding OCT4 protein)(Figure 2H and S2I). Notably, *Nanog* expression was not reduced when cells were treated with dTAG-13 for 72 hours in the naïve state, in agreement with previous studies (Amleh et al. 2009; Williams et al. 2015) (Figure S2J). These results suggest that NELFB and Pol II pausing mediate fine-tuning of gene-regulatory networks during pluripotency transitions rather than the steady-state pluripotent state, but is are dispensable for the induction of formative state transition upon FGF + ACTIVIN treatment. Indeed, pre-treating naïve cells with dTAG-13 for 30 mins, followed by addition of FGF + ACTIVIN for 30 minutes did not affect the induction of immediate FGF targets, such as *Fos* and *Dusp1* (Figure S2K).

### NELF marks active promoters and enhancers in mESCs

To investigate the function of pausing during pluripotency transitions, we first determined the chromatin occupancy of the NELF complex. We performed ChIP-seq of NELFB, NELFE, and SPT5 in *Nelfb^deg^* mESCs maintained in serum/LIF conditions. NELFB and NELFE are components of the NELF complex and are expected to be present solely at Pol II pausing sites, while Spt5 plays an important role in Pol II pausing as well as productive elongation upon phosphorylation by CDK9, making it detectable at both pausing and productive elongation regions (Chen et al. 2018). NELFB, NELFE and SPT5 showed correlated signals at protein-coding gene transcription start sites (TSSs)(Figure 3A, S3A and S3G). NELFB and NELFE peaks (p. adj < 0.05) highly overlapped, suggesting that our NELFB-degron protein fusion maintained its normal chromatin binding capacity (Figure S3B). Annotation of NELFE and NELFB peaks revealed that a subset of called peaks (∼25%) did not correspond to gene TSSs, but instead mapped to intronic and intergenic regions (Figure S3E). We hypothesized that active, transcribed enhancers may show NELF binding in mESCs, and that these likely represented the ∼25% of peaks not associated with gene promoters. Indeed, a large proportion of these peaks mapped to known mESCs enhancers identified in previous studies, and correlated with SPT5 occupancy (Figure 3D, 3E, 3F and S3F)(Whyte et al. 2013). Notably, nearly all super-enhancers contained NELF peaks (Figure 3E). Since super-enhancers have overall higher levels of transcription that typical-enhancers, we suspect that NELF peaks correlate with transcription levels at enhancers (Henriques et al. 2018). These data are in agreement with reports suggesting that Pol II pausing is widespread at enhancers, and suggest that similar to gene TSSs, NELF is a component of the pausing complex at enhancers in mammalian cells (Henriques et al. 2018; Core et al. 2012). Notably, the identification of NELF at enhancers as well as promoters in our system reveals a potential role for enhancer regulation in Pol II pausing/transcription during pluripotent state transitions.

**Figure 3.**
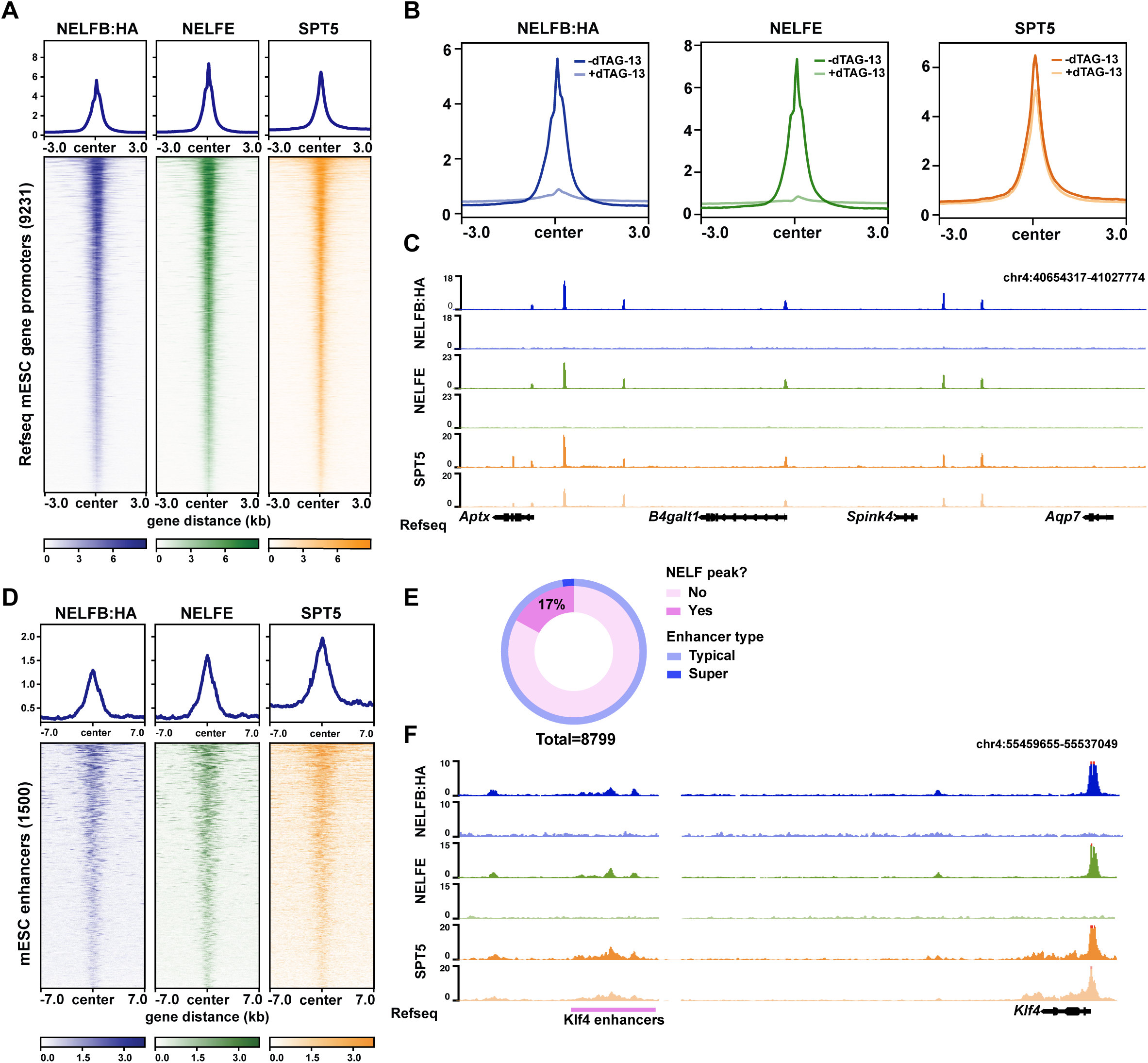
NELF displays widespread binding at promoters and enhancers and *Nelfb^deg^* enables acute clearance of the NELF complex from chromatin. (A) Heatmap of NELFB, NELFE, and SPT5 ChIP-seq signal at active protein-coding genes’ promoters in mESCs. Active promoters were designated as TSSs that contain an SPT5 peak (q. value < 0.05). (B) Metaplot of ChIP-seq signals at promoters defined in (A) with and without 30 mins of dTAG-13. (C) Genome browser shot of a representative region for metaplots in (B). (D) Heatmap of NELFB, NELFE, and SPT5 ChIP-seq signal at mESC-specific enhancers (Whyte et al. 2013). Enhancers with NELF peaks (q. value < 0.05) are shown. (E) Ratio of enhancers and super-enhancers that contain NELF peaks. (F) Genome browser shot of a representative enhancer region showing NELF peaks.

### NELFB deletion results in acute clearance of the complex from chromatin

To test the immediate effect of degrading NELFB on the NELF complex and SPT5, we performed ChIP-seq in matched samples after 30 mins of mESC culture in the presence of dTAG-13. As expected, NELFB peaks were abolished (Figure 3B, 3C, S3A, and S3D). Consistent with the interdependence of individual NELF complex subunits, NELFE peaks were similarly abolished (Figure 3B, 3C, S3A and S3D). Spike-in normalized SPT5 peaks around TSSs showed a global reduction ∼25% (Figure 3B, 3C and S3A). The reduced SPT5 signal suggests that acute disruption of NELF perturbs Pol II pausing but does not abolish transcription entirely. Overall, these results show that *Nelfb^deg^* mESCs can rapidly and specifically remove NELF from chromatin with dTAG treatment and can be used to study the consequences of an acute loss of Pol II pausing. Our results are consistent with recent experiments degrading NELFCD in a human DLD-1 cell line (Aoi et al. 2020).

### NELFB stabilizes Pol II pausing and transcription at promoters and enhancers

Our observations prompted us to study changes in nascent transcription globally upon NELFB depletion in *Nelfb^deg^* mESCs. To assess nascent transcription, we used precision run-on sequencing (PRO-seq)(Kwak et al. 2013; Mahat et al. 2016). PRO-seq identifies the position of transcriptionally engaged RNA polymerases at approximately single base resolution, and allows an assessment of transcription at TSSs, gene bodies, and regulatory elements, including enhancers (Wissink et al. 2019). We were particularly interested in identifying the immediate, direct effects of NELFB loss on transcription. To do so, we treated mESCs in serum/LIF with dTAG-13 for 30- and 60-mins, then collected nuclei for analysis (Figure 4A). We collected 2-3 replicates per condition and used a spike-in to normalize for general transcriptional changes. Replicates showed good correlation (Table S1). Metagene plots revealed a loss of signal at TSSs and gene bodies at both 30- and 60-mins time points (Figure 4B).

**Figure 4.**
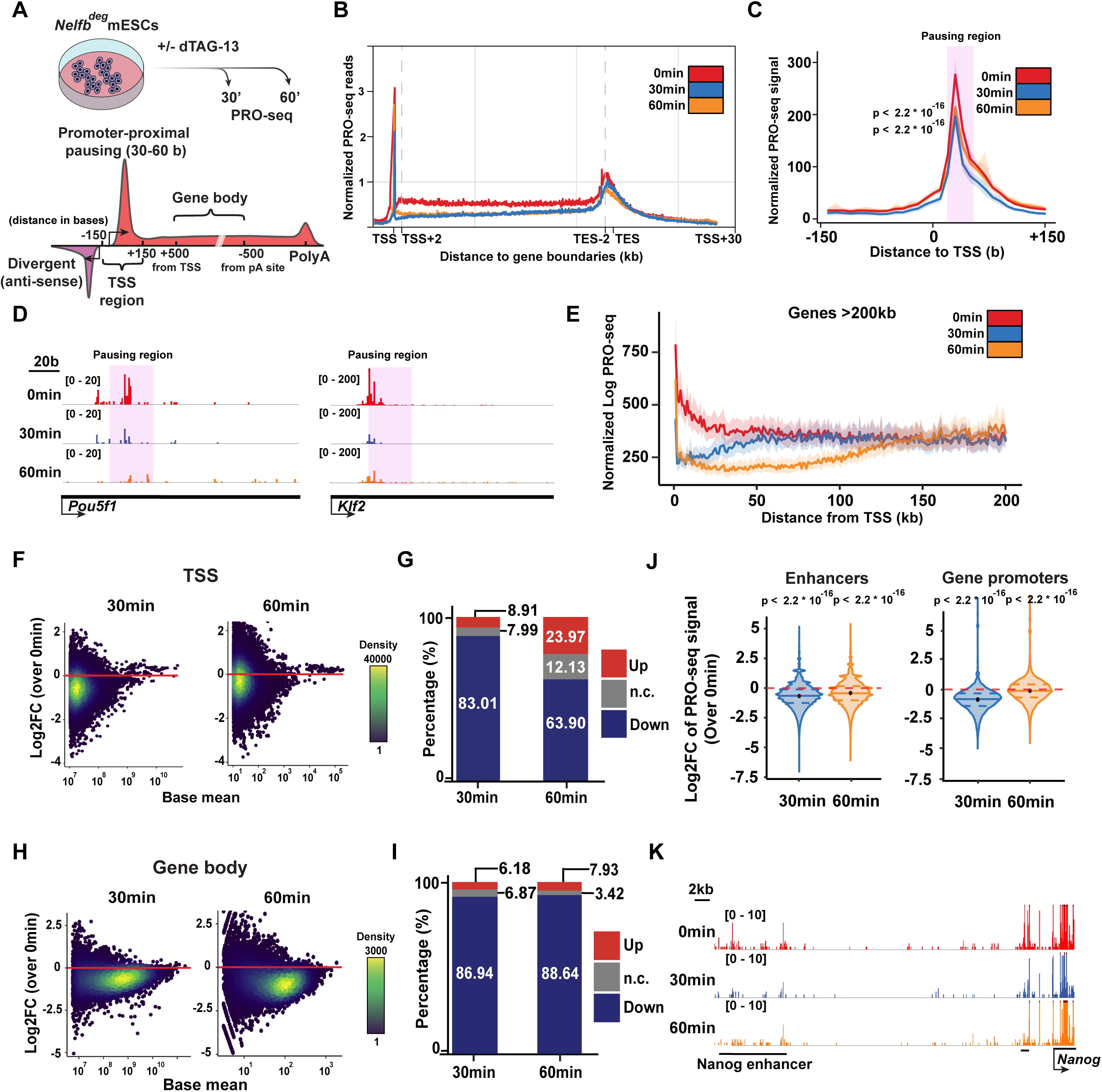
NELF stabilizes Pol II pausing and transcription at promoters and enhancers. (A) (Top)Schematic of treatments of 30 and 60 mins before PRO-seq analysis, and (Bottom) regions of each defined DNA element in following analysis. (B) Metaplot of scaled protein-coding genes’ PRO-seq signal relative to TSS and TES. (C) Metaplot of PRO-seq signal at TSSs. Highlighted region marks proximal-pausing region. Statistical testing was performed using Wilcoxon and paired t-test with similar results. (D) Genome browser shot of TSS regions of example pluripotency genes. Highlighted region marks proximal-pausing region. (E) Metaplot of PRO-seq signal at genes longer than 200kb. (F) Log2 fold change of PRO-seq signal at TSSs calculated using DEseq2. (G) Bar plot showing percentage of up, down, and not changed loci in (F). p. adj. of 0.05 was used as a cutoff. (H) Log2 fold change of PRO-seq signal at gene bodies calculated using DEseq2. (I) Bar plot showing percentage of up, down, and not changed loci in (H). p. adj. of 0.05 was used as a cutoff. (J) Violin plot of TSS Log2 fold change data in figure (F) separated by enhancer vs. protein-coding gene TSSs. Plots show mean, 25^th^, and 75^th^ percentile inside each violin plot. Statistical testing was performed using Wilcoxon and paired t-test with similar results. (K) Genome browser shot of example enhancer signal across treatments

To investigate these changes further, we focused on TSSs. We used published mESCs START-seq data to define the exact positions of TSSs at both gene promoters and regulatory elements (Henriques et al. 2018). TSSs meta-profiles revealed the expected Pol II pause peak 30-50 bases downstream from the TSS (Figure 4C). The peak was significantly and globally reduced when NELFB was depleted (Figure 4C and 4D). Importantly, we identified a drop of PRO-seq signal on gene bodies which extended from TSSs and corresponded with each treatment time and an elongating Pol II speed of ∼1-2kb/min; a drop across the first ∼40kbs of gene bodies in the 30 min treatment group (Figure 4E, S4A and S4C). This phenomenon, previously termed clearing waves, is a result of having elongating polymerases that escape the TSS region prior to dTAG-13 treatment. The presence of clearing waves point to the negative effect of NELFB degradation on transcription which primarily terminates Pol II around the TSS, such that a drop could be seen along gene bodies corresponding to length of treatment time. These results suggest that NELF acts on polymerases close to the TSS to enable efficient transition of polymerases from pausing to productive elongation. Our findings place NELF as a positive effector required for transcription to proceed effectively and highlights the power and specificity of our degron system.

To determine signal changes at each locus in a pair-wise manner, we assessed differential expression at all active TSSs in mESCs and gene bodies using DEseq2 (Love et al. 2014). In agreement with our previous observations, we noted a global reduction in transcription at TSSs, on average, within 30 min (Figure 4F-J, S4B and S4D). Notably, the reduction at 60 min was conserved at enhancer TSSs and gene bodies, but not at gene TSSs (Figure 4J). This recovery of transcription at gene TSSs from 30 min to 60 min was not found in the canonical Pol II pausing region (∼30 bases from TSSs), but further downstream in an apparent redistribution of the pause peak in the absence of NELF which presumably stabilizes the pause-position to 30-50 bases from TSSs (Figure 4C and 4D)(Aoi et al. 2020).

To define the properties of the promoters that displayed Pol II redistribution, we selected a list of significantly recovering gene TSSs (404 genes; at 30 mins: down p. adj < 0.05; at 60 min: up p. adj < 0.05) and measured levels of NELF and the active promoter H3K4me3 mark. Additionally, we inferred the rates of initiation and pause-release at these promoters using a recently described statistical model (see methods)(Siepel 2021). Of note, the initiation and release rates model has been developed to function under steady-state conditions without perturbation. The rates calculated are relative and do not reflect absolute numbers of initiation or release events, which enables intra-sample comparison only. We found that genes exhibiting a redistribution of Pol II consistently harbored high signals for NELFB, NELFE, and H3K4me3, suggesting that these are highly active promoters with significant occurrence of pausing (Figure S4E and S4F). Measuring the initiation and pause-release rates showed that these promoters have a higher initiation rate and a lower release-rate, indicating that these genes may have high initiation rates whereas the rate of pause release is rate limiting to transcriptional activation (Figure S4G). Transient transcriptome sequencing (TT-seq) can detect nascent transcription as well as terminated transcripts, making it able to measure initiation rates experimentally (Schwalb et al. 2016). Measuring the signal of recovering genes in publicly available mESCs TT-seq data confirmed that these promoters are more active in untreated mESCs, indicating higher initiation rates (Figure S4H)(Shao et al. 2021). Overall, we found that Pol II pausing correlates with transcriptional activity globally, and that NELF plays a specific role in stabilizing paused polymerases at a defined position 30-50 bases downstream of TSSs which enables efficient transition from initiation to productive elongation.

### Pol II pausing balances induced and repressed gene regulatory networks during pluripotency transitions

Having established the validity of the *Nelfb^deg^* mESCs and acute molecular consequences of NELFB depletion on Pol II pausing and transcription, we sought to analyze the effect of depleting NELF during pluripotent state transitions. We opted to use a transient pulsed NELF degradation approach in *Nelfb^deg^* mESCs. As described earlier, this treatment regimen was able to recapitulate the state transition defects observed in embryos, while minimizing secondary effects. This experimental design enables us to assess how a minimal perturbation of Pol II pausing during transitions would affect transcription of transitioning cells. Transitioning cells were treated with dTAG-13 for 1 hour between 23-24 hours of the transitioning protocol in FGF + ACTIVIN containing medium, followed by washing and continued culture in the presence of FGF + ACTIVIN, but in the absence of dTAG-13 for total of 72 hours (Figure 5A). This treatment results in acute depletion of NELFB at the 24 hour time-point, and a recovery over the following 24 hours, as we have shown previously (Figure S2D). Samples were collected for PRO-seq at 24-, 28-, 48- and 72 hours. The first two time-points represent intermediate points of the transitions, while the latter two represent fully transitioned EpiLCs/formative states. Our analysis focused on pluripotency associated genes that are either differentially expressed or genes that maintain a comparative level of expression during state transitions. We identified genes that were upregulated, downregulated, and shared between the naïve (0 hour), and formative (48hour), stages using DEseq2 in untreated samples (-2.5 > Log2FC > 2.5, p. adj < 0.05). These groups included many expected genes that are specific to, or are shared between states, thereby validating the transition of these cells; up: *Otx2, Pou3f1, Fgf5, Fgf15*; down: *Klf4, Klf2, Nr0b1, Nanog*; shared: *Sox2, Pou5f1* (Figure 5B).

**Figure 5.**
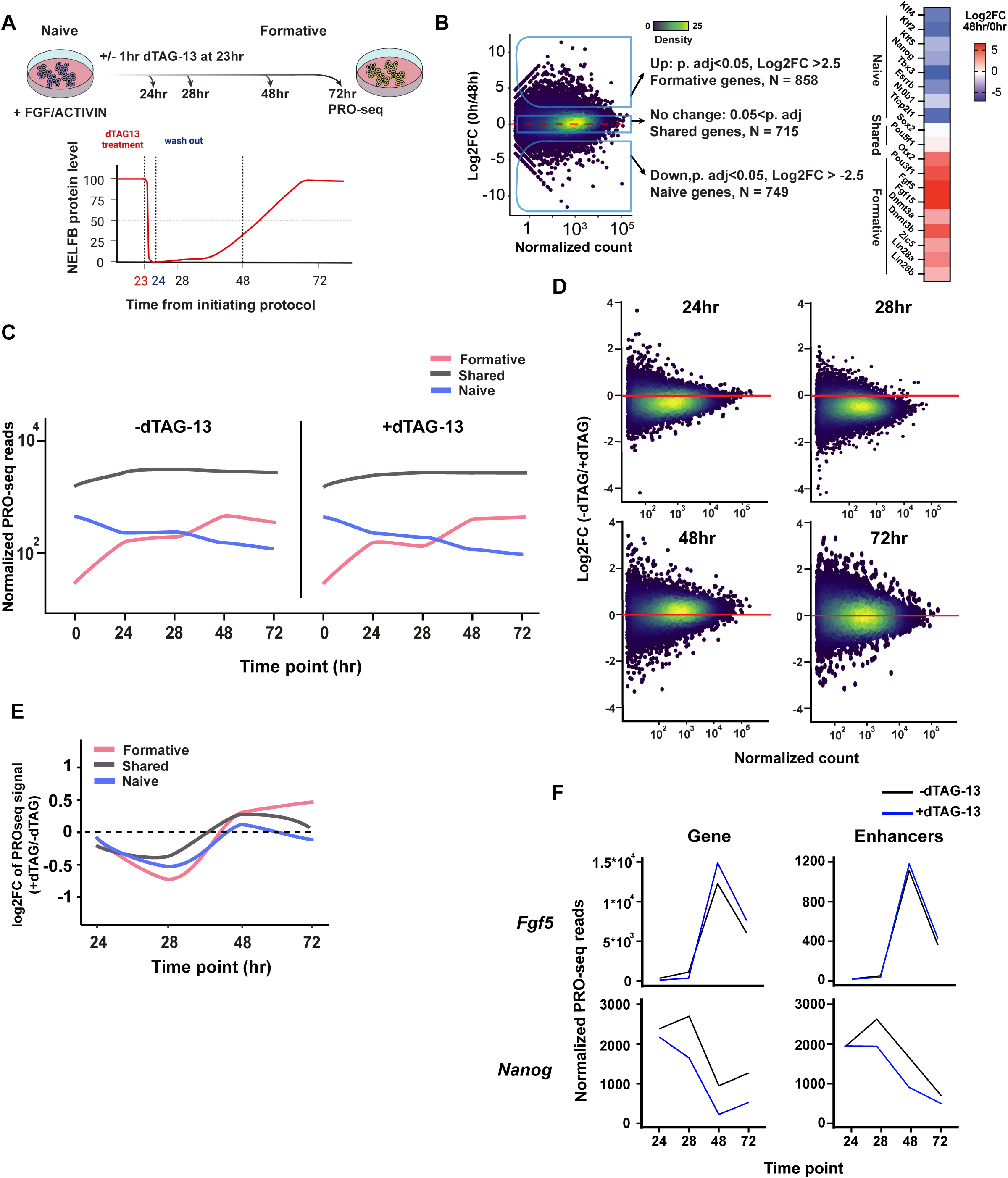
NELF balances gene induction and repression during pluripotency transitions. (A) (Top) Schematic of experiment and analysis timepoints. (Bottom) Schematic of NELFB protein levels during the experiment following transient depletion. (B) (Left) Log2 fold change of PRO-seq data gene expression between 0hr and 48hr which was used to define naïve genes, formative genes, and shared genes. (Right)Heatmap of Log2 fold change of known naïve and formative markers. (C) Mean normalized PRO-seq reads per gene in each gene class during the transition. Full data range is shown in Figure S5A and heatmaps in Figure S5B. (D) Log2 fold change of PRO-seq data gene expression at each time point of the analysis using DEseq2. (E) Mean log2 fold change of PRO-seq data gene expression at each time point of the analysis per gene group. Full data range is shown in Figure S5C. (F) Normalized gene expression/reads from PRO-seq data at candidate genes and their associated enhancers during the transition protocol. Other genes shown in Figure S5F.

To assess whether NELFB depletion during transitions influenced the cells’ ability to initiate transitions, we generated metaplots and heatmaps of naïve and formative genes at each timepoint with and without dTAG-13 treatment (Figure 5C, S5A, and S5B). The general trend suggested that treated cells maintained expression of the same genes as untreated cells in agreement with the ability of *Nelfb^deg^* mESCs to induce formative markers when cultured in dTAG-13 and *Nelfb^-/-^* embryos upregulating formative epiblast markers.

To quantify these observations, we performed a pair-wise differential expression analysis using DEseq2 and tracked the trend of each gene group during the transition. In agreement with our previous results, we found that the 24 and 28 hour time-points showed a global decrease in transcription when compared to non-treated timepoint-matched controls (Figure 5D). This global decrease is largely recovered in the terminal points at 48 and 72 hours, most likely due to recovery of NELFB protein (Figure 5D). To directly determine the effect of NELFB depletion on induced (formative-specific), repressed (naive-specific), and shared genes between both states during the transition, we compared the change in expression of these groups of genes at each timepoint with and without dTAG-13. While all groups show initial down regulation, state-specific groups (naïve and formative) were more severely affected (Figure 5E and S5C). By 72 hours, shared genes between naive and formative states showed minimal change, while genes induced as cells entered the formative state show a stronger induction, and genes repressed in the naïve state showed stronger silencing, with several candidate genes showing this trend (Figure 5E, 5F, and S5F). These data offer evidence for an involvement of Pol II pausing in mediating the levels of expression of genes which are either up or down regulated during pluripotency transitions (Figure 5E and S5C). In the absence of pausing, gene activation and repression are mis-regulated during pluripotent state transitions.

Previous studies have linked enhancer transcription to target gene promoter activity (Hah et al. 2013; Kim et al. 2010). Given that NELF and Pol II pausing can occur at enhancers, we wanted to assess the enhancer landscape during pluripotent state transitions. To do so, we employed two approaches. First, we used the dREG algorithm to identify transcriptional regulatory elements (TREs), genomic regions that have putative roles in gene regulation at the formative stage (Wang et al. 2019). Overall, TREs showed downregulated expression in samples treated with dTAG-13 at most timepoints, reemphasizing the role of NELF and pausing in maintaining enhancer activity (Figure S5D). To identify specific changes at putative enhancers for genes of interest, we selected TREs that fall within a topologically associated domain (TAD) of a gene of interest and were marked by H3K27ac histone modifications for active enhancers. This strategy enabled us to identify several high confidence putative enhancers for genes (Figure S5E; see methods). We applied this approach to the *Nanog* and *Fgf5* loci as representative genes that are repressed and induced, respectively during the naive to a formative state transition. We find that *Nanog* and *Fgf5* enhancer activity mirrored the trend in their respective gene expression (Figure 5F). The observed changes are consistent with the presence of Pol II pausing at enhancers, and the coupling between transcription at enhancers and associated target genes. Overall, these results detail the effects of perturbing pausing during pluripotency transitions at the transcriptional level, where Pol II pausing plays a role in balancing genes and enhancers’ induction and repression during state transitions.

### NELF recruitment to chromatin is enhanced during pluripotency transitions

Previous studies on the function of Pol II pausing and NELF in mESCs suggested that Pol II pausing is not required to maintain pluripotency (Amleh et al. 2009; Williams et al. 2015). Our results in embryos and mESCs using controlled NELFB depletion support these studies and extend them by suggesting that Pol II pausing plays a key role during changes in cell states. We hypothesized that if this is the case, *de novo* recruitment of NELF to chromatin may be observed during pluripotency transitions. To test this hypothesis, we measured NELFB levels in the chromatin fraction during the transitioning period. In support of our results, we found a significant increase in chromatin bound NELFB but not in whole cell lysates observed at 24 and 48 hours of transitioning in FGF + ACTIVIN (Figure 6A, 6B and S6A). Notably, this increase was not observed at 4 hours of transitioning, suggesting that NELFB recruitment is not initiated during the acute phase of FGF + ACTIVIN growth factor stimulation, but rather during the rewiring of transcriptional networks that follow.

**Figure 6.**
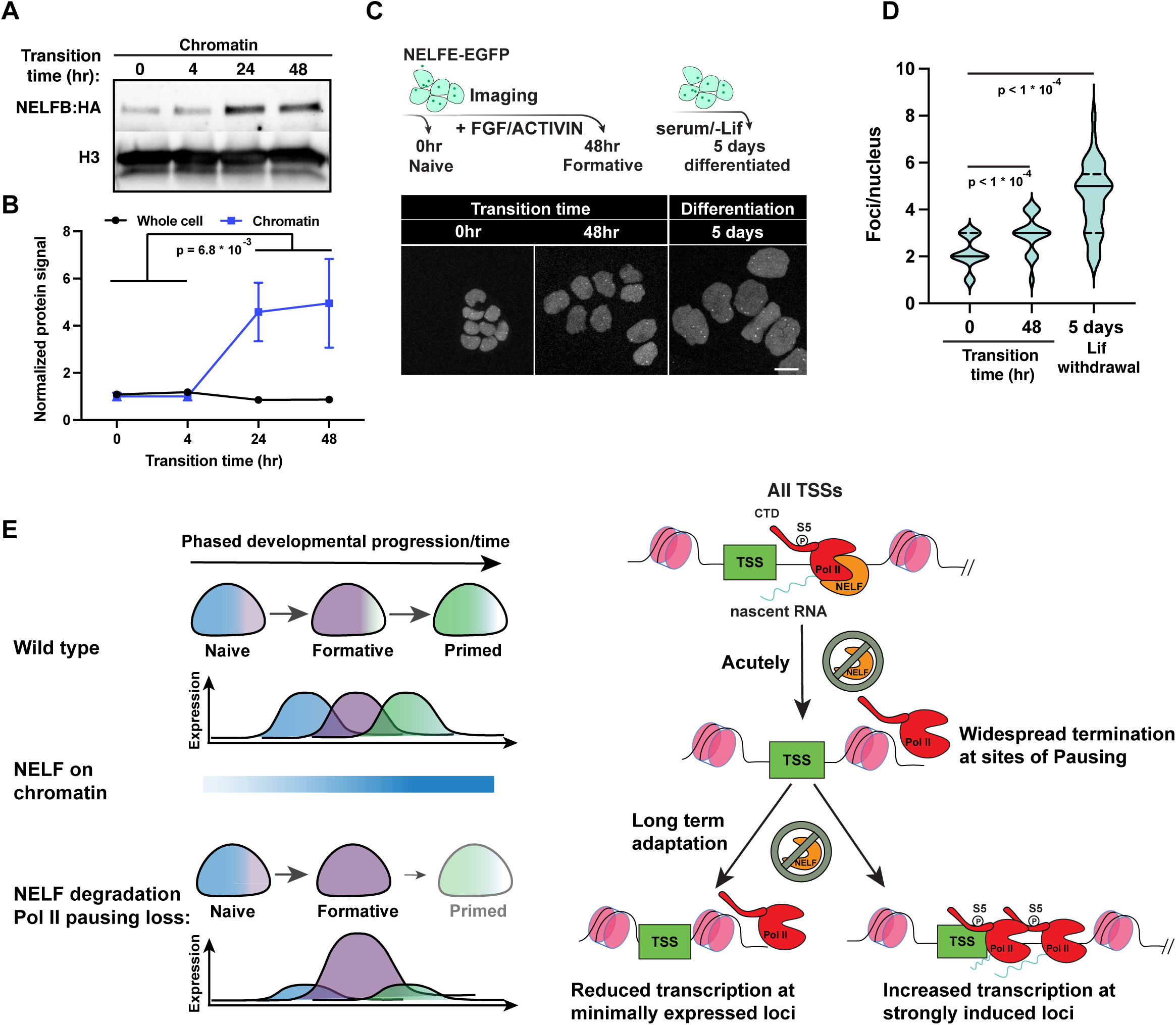
NELF is recruited to chromatin during pluripotency transitions. (A) Chromatin fraction western blot of cells during pluripotency transitions. (B) Quantification of NELFB in chromatin and whole cell lysates during pluripotency transitions. Statistical testing was performed using a t-test. Each time point includes two biological replicates. (C) Imaging of NELFE-EGFP in naïve, formative, and randomly differentiated mESCs. Top, schematic of the experiment. Bottom, images of select time points. (D) Violin plot of number of NELF bodies per nuclei in conditions presented in (C). Statistical testing was performed using a t-test. Mean, 25^th^, and 75^th^ percentiles are shown inside each violin plot. (E) Schematic of proposed NELF/Pol II pausing function during pluripotency transitions at the molecular and cellular levels.

To extend these observations, we took advantage of work that identified a putatively liquid-liquid phase separated compartment, referred to as NELF bodies, as sites of NELF-mediated transcriptional regulation (Narita et al., 2007; Rawat et al., 2021). To visualize NELF bodies, we generated a clonal transgenic NELFE-EGFP fusion on our *Nelfb^deg^* background mESC line (Figure S6B). In dTAG-13 untreated conditions, distinct foci (∼2-4 per nucleus), could be visualized, consistent with previous observations. However, NELFB depletion resulted in complete dissolution of NELF bodies without affecting overall fluorescence levels, further demonstrating an interdependence between the subunits of the NELF complex, and suggesting that these bodies represent hubs of transcriptional regulation (Figure S6C-D). We hypothesized that cells would display a greater number of NELF bodies during periods of transition, for example when transitioning pluripotent states, or changing their fate, as opposed to steady state conditions. Indeed, we found a significant increase in the number of NELF bodies per nucleus upon pluripotency transition, as well as during the differentiation of mESCs maintained in serum containing medium in the absence of LIF (Figure 6C, 6D, S6D and S6E). Our results suggest that *de novo* NELF recruitment to chromatin occurs during pluripotency transitions, presumably to attenuate and buffer gene induction and repression to ensure a smooth transition between sequential cell states.

## DISCUSSION

The discovery of Pol II pausing at heat shock genes represented an additional layer of gene regulation (Rougvie and Lis 1988). Subsequent work defined the protein complexes involved in this step, including the NELF and DRB sensitivity inducing factor, DSIF (SPT5 and SPT4), as major regulators of Pol II pausing (Wu et al. 2003; Gilchrist et al. 2012). Further work demonstrated that Pol II pausing occurs globally in metazoans and can regulate the transcriptional output of a variety of signaling pathways (Abuhashem et al. 2022; Danko et al. 2013; Liu et al. 2015; Nechaev et al. 2010). Recent structural studies have provided high-resolution maps of the paused Pol II complex showing how NELF and unphosphorylated SPT5 can block elongation of Pol II and sterically inhibit the formation of new pre-initiation complexes (PICs), confirming the potential of Pol II pausing to act as a bottleneck step in transcription (Vos et al. 2018a, 2018b).

Previous studies identified roles of Pol II pausing in cultured cells, as well as model organisms such as *Drosophila*, Zebrafish, and mice. These roles revolved around modulating responses to several signaling pathways. In mouse, NELF was found to have an essential role in embryonic development and for enabling the differentiation of mESCs in culture via regulating FGF signaling (Amleh et al. 2009; Williams et al. 2015). These studies relied on long-term genetic knockout or siRNA approaches which result in secondary defects that may mask primary and acute functions of NELF. Here, we sought to understand the direct function of Pol II pausing in early mammalian development by applying acute protein depletion to interrogate the molecular and temporal requirements of Pol II pausing *in vitro* in mouse ESCs, in parallel with studies *in vivo* in mutant embryos. We identify state transitions within the pluripotent epiblast of the embryo, and modelled by pluripotent stem cells in culture, preceding the onset of germ layer differentiation, as a key process that requires Pol II pausing to achieve smooth state transitions, and ultimately differentiation of pluripotent cells.

The timing of the defect characterized in mouse post-implantation embryos is consistent with previous studies identifying the role of *Nelfb* in mESC differentiation *in vitro* (Amleh et al. 2009; Williams et al. 2015). It is, however, notable that initial cell fate specification events in the blastocyst and peri-implantation stages were unaffected in the absence of NELFB. As pluripotent cells progress from their initial naive to a later primed state, they prepare to exit pluripotency in favor of germ layer specification and differentiation. Pluripotent cells therefore need to calibrate gene expression for precise spatiotemporal control of cellular differentiation. Our data suggests that Pol II pausing mediates cell state transitions by balancing gene regulatory networks during transitions. This model is supported by previous studies at the molecular and cellular levels. Molecularly, profiling of Pol II pausing across pre-implantation mouse development has identified a distinct reduction in Pol II pausing following Zygotic Genome Activation (ZGA) until late blastocyst stage, at which point it is re-established (Liu et al., 2020). We recently showed that NELF is required at this specific stage prior to ZGA to regulate the major ZGA wave in mouse embryos (Abuhashem and Hadjantonakis 2021). At the cellular level, several studies investigating tissue-specific *Nelfb* knockouts have revealed that functional defects are observed when *Nelfb^-/-^* tissues are challenged by an external stimulus, such as an injury or an infection or the need to regenerate in the context of muscle stem cells, the uterine and intestinal walls, and in macrophages (Hewitt et al. 2019; Ou et al. 2021; Robinson et al. 2021; Yu et al. 2020). Our model, suggesting that Pol II pausing acts to fine-tune transcription during state transitions, explains the defects observed in both the present and previous studies.

Leveraging the dTAG system to acutely deplete NELFB at specific time-points allowed us to address why Pol II pausing may be particularly important during pluripotent state transitions and, potentially, in other contexts where cells transition between different states. By combining the fine temporal control of protein expression with the resolution of PRO-seq data, we were able to assess the direct effects of NELFB depletion on global transcriptional activity while bypassing the secondary effects of disruption Pol II pausing on cell proliferation. Our data suggest that disrupting Pol II pausing as cells transition between successive states results in dysregulation of induced and repressed genes and their enhancers. Specifically, Pol II pausing appears to limit the induction of gene networks and delay the loss of repressed gene networks. Super-induction of state specific genes in the absence of NELF, as observed in our data, effectively functions as an overexpression of state-specific genes, limiting the ability of cells to exit the current state and acquire the subsequent (Figure 6E). This conclusion is supported by the observation of increased chromatin recruitment of NELF during pluripotency transitions. Notably, a similar loss of Pol II pausing at the earliest stage of state transitioning, hour 0, did not result in a defect, and we did not observe increased NELF chromatin recruitment at this stage, 0 – 4 hours of transitioning. These data suggest that Pol II pausing is not necessarily required for acute responses to the cytokines used here to drive pluripotent state transitions, FGF and NODAL, and potentially other signals. This is in line with normal induction of early-release genes, such as *Fos*, after NELFB degradation and a recent analysis of FGF signaling in mESCs concluding that Pol II recruitment, rather than release, is the rate limiting step in the activity of FGF/ERK signaling pathway (Hamilton et al. 2019).

Super-induction of highly active loci in the absence of Pol II pausing has been observed previously, and molecularly may be due to Pol II pausing acting as a rate limiting step at highly active loci (Henriques et al. 2018; Yu et al. 2020). Conversely, loss of minimally expressed/repressed genes could result from increased nucleosome occupancy in the absence of a paused Pol II (Figure 6E)(Gilchrist et al. 2010; Henriques et al. 2018). Indeed, we could observe both effects at the same locus, *Nanog*, depending on its expression status, further supporting a link between Pol II pausing role and the level of gene expression. Importantly, our analysis does not refute previous results suggesting that FGF signaling is attenuated in *Nelfb^-/-^* mESCs, but rather suggests that these defects are most likely secondary (Williams et al. 2015).

Our data suggest that NELF-enforced Pol II pausing is widespread at enhancers and promoters. Depleting NELF destabilizes and terminates paused transcripts. These observations highlight a general positive effect of NELF-enforced Pol II pausing on transcription. The presence of paused Pol II can regulate and limit transcription from a certain locus, however, its loss results in destabilizing this important regulatory step and not in release of productive elongating polymerases. Furthermore, at gene promoters that have high initiation rates, NELF centers the paused polymerase 30-50 bases downstream of the TSS, and upon its depletion, polymerases extend further downstream, but do not produce productive elongation. These observations are consistent with a study performing acute depletion of NELFCD, which resulted in the formation of a “second pause” position of promoter-proximal Pol II (Aoi et al., 2020).

In summary, by performing comprehensive investigation of Pol II pausing function in a relevant developmental context, and comparative *in vivo* (embryo) and *in vitro* (mESCs) models, we propose a model whereby pausing functions as a rheostat for changing transcriptomes during cell state transitions.

## MATERIAL AND METHODS

### Materials availability

Request for reagents should be directed to and will be fulfilled by the lead contact, Anna-Katerina Hadjantonakis

### Cell lines

ATCC E14 ES cell line was cultured on 0.1% gelatin (Millipore) coated tissue-culture grade plates in a humidified 37°C incubator with 5% CO_2_ (Kiyonari et al., 2010). For routine culture, cells were grown in Serum/LIF conditions: DMEM (Gibco), supplemented with 2 mM L-glutamine (Gibco), 1x MEM non-essential amino acids (Gibco), 1 mM sodium pyruvate (Gibco), 100 U/ml penicillin and 100 U/ml streptomycin (Gibco), 0.1 mM 2-mercaptoethanol (Gibco), 15% Fetal Bovine Serum (Gibco), and 1000 U/ml of recombinant leukemia inhibitory factor (LIF).

To model different stages of pluripotency, cells were initially cultured in N2B27 + 2i/LIF for 4 days to induce naïve pluripotency, equivalent 0 hr in this study. N2B27 comprised of 50% Neurobasal medium (Gibco) with 100x N2 supplement (Gibco), 50% DMEM/F12 (Gibco) with 50x B27 supplement (Gibco), 2 mM L-glutamine (Gibco), 100 U/ml penicillin and 100 U/ml streptomycin (Gibco), 0.1 mM 2-mercaptoethanol (Gibco), 1% KnockOut Serum Replacement (Gibco). To initiate transitions, we followed the EpiLC conversion protocol. Plates were coated with 16 μg/ml of Fibronectin (Millipore) in PBS for 30 mins at 37°C, followed by two washes of PBS. Naïve cells were plated at 25*10^3^/cm^2^ in N2B27 supplemented with 12 ng/ml FGF2 and 20 ng/ml ACTIVIN A (Peprotech). Medium changes were done daily for all conditions.

### Plasmid generation

Three plasmids were generated for this study: (1) Cas9 vector to target the C-terminus of *Nelfb* gene. PX459 vector (addgene #62988) was digested using BbsI-HF (NEB) and single guide RNA targeting *Nelfb* was annealed (Ran et al., 2013), (2) Homology directed repair (HDR) vector containing the insert FKBP^F36V^ tag, 2x HA tag, self-cleaving P2A sequence, and Puromycin resistance, flanked by 1 kb *Nelfb* HDR sequences. The insert was obtained from pCRIS-PITCHv2-dTAG-BSD (addgene #91795)(Nabet et al., 2018). The plasmid backbone (pBluescript), *Nelfb* HDR sequences, and the insert were amplified using Q5 polymerase (NEB) and the plasmid was constructed using NEBuilder HiFi DNA assembly (NEB). (3) *Nelfe*-EGFP vector as a fluorescent reporter of NELF bodies. *Nelfe* cDNA was amplified using Q5 polymerase (NEB). Linker-EGFP and PGK backbone were amplified from pHaloTag-EGFP (addgene #86629) and PGKneobpa (addgene #13442) respectively. plasmid was constructed using NEBuilder HiFi DNA assembly (NEB).

### Genome editing

To generate *Nelfb^deg^* mESCs, 3 million cells were transfected with 10ug PX459-Nelfb_sgRNA and 10ug Nelfb_left-FKBP^F36V^- 2xHA- P2A-BSD-Nelfb_right. Cells were transfected using Lonza P3 Primary Cell 4D-Nucleofector^TM^ X 100 ul cuvettes (Lonza). Following transfection, cells were plated on a 10 cm dish (Falcon) coated with MEFs. 48 hrs post transfection, correctly targeted cells were selected for using 6 ug/ml Blasticidin (InvivoGen) for 5 days. Surviving cells were split 1000 cells/10 cm dish and maintained for 9 days under Puromycin selection. Surviving clones were picked under a stereomicroscope, expanded, and genotyped for the insert.

### Mouse strains and husbandry

All animal work was approved by MSKCC Institutional Animal Care and Use Committee (IACUC). Animals were housed in a pathogen free-facility under a 12-hr light cycle. Mouse strains used in this study were *Nelfb^+/-^* and wild-type CD-1/ICR (Charles River). *Nelfb^+/-^* mice were imported from Karen Adelman lab (Jax #033115). The imported mice had a floxed allele. Following crossing with Zp3-cre (Jax #003651), heterozygous knockout progeny was identified and expanded.

### Cells dTAG treatment

dTAG-13 (Bio-Techne) was reconstituted in DMSO (Sigma) at 5 mM. dTAG-13 was diluted in maintenance medium to 500 nM and added to cells with medium changes for the specified amounts of time.

### Embryo collection

For all experiments, embryos were obtained via natural mating of 6-12 weeks of age females with 7 – 16 weeks of age males. For preimplantation stages, embryos were recovered by flushing the uterine horns (E3.25 – E4.5). These dissections were carried out in flushing and holding medium (FHM, Millipore) as described (Behringer et al., 2014).

For post-implantation embryos (E5.5 – E7.5), the uterine horns were retrieved and cut into single decidual swellings in 5% Newborn Calf Serum in DMEM/F12 (Gibco). Embryos were dissected out by removing the uterus wall and decidual tissue. The parietal endoderm was removed carefully with the ectoplacental cone.

### Immunofluorescence

For cultured mESCs, cells were plated on u-Slide 8 well (ibidi), washed with PBS+/+ and fixed in 4% PFA (electron microscopy sciences) in PBS+/+ for 10 min at room temperature. Fixed cells were washed two times with PBS+/+, one time with wash buffer; 0.1% Triton X-100 (Sigma) in PBS+/+, then permeabilized in 0.5% Triton X-100 (Sigma) in PBS+/+ for 10 min. Cells were then blocked with 3% Donkey Serum (Sigma) and 1% BSA (Sigma) for 1 hr at room temperature. Cells were then incubated with primary antibodies in blocking buffer at 4°C over night (antibodies and concentrations in Table S1). Cells were then washed three times in wash buffer, and incubated with suitable donkey Alexa Fluors^TM^ (Invitrogen, 1:500) for 1 hr at room temperature. Cells were then washed three time wish wash buffer, the last containing 5 μg/ml Hoechst 33342 (Invitrogen), then imaged.

For E3.25-E4.5 pre-implantation embryos, the zona pellucida was removed by incubation in acid Tyrode’s solution (Sigma) at 37°C for 2 min. Embryos were subsequently washed briefly in PBS+/+ before fixation in 4% PFA for 10 mins at room temperature. Fixed embryos were washed in 0.1% Triton X-100 in PBS+/+ (PBX)for 5 min, permeabilized in 0.5% Triton X-100 (Sigma) in PBS+/+ for 5 min, washed again for 5 min in PBX, and blocked in 2% horse serum (Sigma) in PBS+/+ for 1 hr at room temperature. Embryos were incubated in primary antibodies diluted in blocking solution over night at 4°C. Embryos were then washed three times for 5 min each in PBX and blocked again for 1 hr at room temperature prior to incubation with secondary antibodies. Secondary antibodies diluted in blocking solution were applied for 1 hr at 4°C. Embryos were then washed twice for 5 min each in PBX and incubated with 5 ug/ml Hoechst 33342 (Invitrogen) in PBS for 5 min or until mounting for imaging. The following primary antibodies were used: goat anti-GATA6 (R&D Systems, 1:100), mouse anti-CDX2 (BioGenex, 1:200), rabbit anti-NANOG (CosmoBio, 1:500). Secondary Alexa Fluor-conjugated antibodies (Invitrogen) were used at a dilution of 1:500. DNA was visualized using Hoechst 33342.

For E6.5 and E 7.5, Embryos were washed briefly in PBS+/+ before fixation in 4% PFA for 20 mins at room temperature. Fixed embryos were washed in 0.1% Triton X-100 in PBS+/+ (PBX) for 5 min, permeabilized in 0.5% Triton X-100 (Sigma) in PBS+/+ for 20 min, washed again for 5 min in PBX, and blocked in 3% horse serum (Sigma) in PBX for 1 hr at room temperature. Embryos were incubated in primary antibodies diluted in blocking solution over night at 4°C. Embryos were then washed three times for 10 min each in PBX and blocked again for 1 hr at room temperature prior to incubation with secondary antibodies. Secondary antibodies diluted in blocking solution were applied over night at 4°C. Embryos were then washed three times for 5 min each in PBX and incubated with 5 ug/ml Hoechst 33342 (Invitrogen) in PBX for 1 hr or until mounting for imaging.

### Image data acquisition

Fixed immunostained samples were imaged on a Zeiss LSM880 laser scanning confocal microscope. Pre-implantation embryos were mounted in microdroplets of 5 μg/ml Hoechst 33342 in PBS+/+ on glass-bottomed dished (MatTek) coated with mineral oil (Sigma). Embryos were imaged along the entire z-axis with 1μm step using an oil-immersion Zeiss EC Plan-Neofluar 40x/NA 1.3 with a 0.17 mm working distance. For post-implantation embryos, a similar setup was used but with an air Plan-Apochromat 20x/NA 0.75 objective. Super resolution imaging of Nelfe-EGFP was done on a Zeiss Elyra 7 with lattice SIM using an oil-immersion Zeiss Plan-Apochromat 63x/NA 1.4 objective.

### Western blotting

For cells, 350ul of lysis buffer; 1x Cell Lysis Buffer (Cell Signaling) with 1mM PMSF (Cell Signaling) and cOmplete^TM^ Ultra protease inhibitor (Sigma), was added to a 90% confluent 6-well dish (Falcone) after washing with PBS-/-. Cells were incubated with lysis buffer for 5 min on ice, then scraped and collected. Samples were sonicated for 15 seconds to complete lysis at, then spun down at 12,000x g for 10 min at 4°C. The supernatant was collected, and protein concentration measured using Pierce^TM^ BCA Protein Assay Kit (Thermo). 10-20 ug of protein was mixed with Blue Loading Buffer (Cell Signaling) and 40 mM DTT (Cell signaling). Samples were boiled at 95°C for 5 min for denaturation. To prepare cellular compartment fractions, Subcellular Protein Fractionation kit was used (Thermo) according to the manufacturer’s instruction.

Samples were run on a BioRad PROTEAN system and transferred using Trans-Blot Semi-Dry Transfer Cell (BioRad) to a nitrocellulose membrane (Cell Signaling) following manufacturer’s instructions and reagents. Membranes were then washed briefly with ddH2O and stained with Ponceau S (Sigma) for 1 min to check for transfer quality, and as a loading control. Membranes were then washed three times with TBST; 0.1% Tween 20 (Fisher) in TBS. Membranes were blocked with 4% BSA in TBST for 1 hr at room temperature and subsequently incubated with primary antibodies diluted in blocking buffer at 4°C over night. They were then washed three times with TBST, then incubated with secondary antibodies in blocking buffer for 1 hr. Washed three times with TBST, incubated with ECL reagent SignalFire^TM^ for 1-2 min and imaged using a ChemiDoc (BioRad). Primary and secondary antibodies are listed in (Table S1) .

### RT-qPCR

RNA was extracted from samples using TRIzol (Thermo) following the manufacturer’s instructions. 1μg of RNA was used to generate cDNA using the QuantiTect reverse transcription kit (Qiagen). qPCR reaction was performed using PowerUp SYBR green mastermix (thermo) and a BioRad CFX96. Used primers are available in (Table S1).

### ChIP-seq

25 million cells were collected for each sample/replicate. Cells were crosslinked in 1% PFA (Electron Microscopy Sciences) in PBS for 10 min at room temperature. Following quenching with 125mM glycine (Sigma) for 5 min at room temperature, cells were washed twice with PBS instructionsthen suspended in lysis buffer: 10□mM Tris pH□8, 1□mM EDTA and 0.5% SDS (Sigma); 20□×□10^6^ cells per 400□μl. To shear chromatin, samples were sonicated using a Bioruptor^®^ Pico sonication device (Diagenode) for 12 cycles, 30 seconds on/30 seconds off then pelleted at the maximum speed for 10□min at 4□°C. The supernatant was diluted five times with dilution buffer: 0.01% SDS, 1.1% Triton X-100, 1.2□mM EDTA, 16.7□mM Tris pH□8 and 167□mM NaCl (Sigma) then incubated with primary antibodies at 4°C over night. Protein G Dynabeads^TM^ (Thermo) were blocked at 4°C over night using 100 ng/10μl of beads. The next day, beads were added to samples at 20 μl per sample for 3 hour at 4°C. Using a magnet to stabilize the beads, they were washed twice in low-salt buffer (0.1% SDS 1% Triton X-100, 2□mM EDTA, 150□mM NaCl and 20□mM Tris pH□8), twice in high-salt buffer (0.1% SDS, 1% Triton X- 100, 2□mM EDTA, 500□mM NaCl and 20□mM Tris pH□8), twice in LiCl buffer (0.25□M LiCl, 1% NP-40, 1% deoxycholic acid, 1□mM EDTA and 10□mM Tris pH□8) and once in TE buffer (10 mM Tris pH 8, 0.1 mM EDTA). Subsequently, the DNA was eluted from the beads by incubating with 150□μl elution buffer (100□mM NaHCO_3_ and 1% SDS) for 20□min at 65□°C with vertexing using Eppendorf ThermoMixer C (Eppendorf). The supernatant was collected, reverse crosslinked by incubation overnight at 65□°C in the presence of proteinase K (Roche), and cleaned by RNase A (Thermo) treatment for 1□hour at 37□°C, the DNA was purified using a DNA clean and concentrate kit (Zymo Research). Spt5 ChIP samples were spiked in with 10% human HEK 293T cells to perform normalized quantification of signal.

### ChIP-seq analysis

Reads were aligned to mm10 and filtered using the following pipeline (https://github.com/soccin/ChIP-seq). Briefly, reads were aligned using Bowtie 2.3.5, then filtered using a MAPQ>30. Properly paired reads were kept. Resulting BAMs were used to generate BigWigs using DeepTools (https://deeptools.readthedocs.io/en/develop/). BigWigs were normalized to 10 million. For SPT5, the samples were aligned to an index with both mm10 and hg38 to normalize to human cells spike-in. The normalization was applied as a scale factor during BigWigs generation, where the scale factor is the multiple required for each spike-in to be equal to the average of all spike-ins. Peak calling was performed using MACS2 and q-value < 0.05. Shared peaks across replicates were analyzed. Downstream analysis was performed in Rstudio 4.1.2 using Bioconductor packages and deepTools to generate heatmaps.

### PRO-seq (sample preparation and library prep)

5-15 x 10^6^ cells were detached using tryspin (Thermo), then resuspended in 500 μl wash buffer: 10 mM Tris-Cl pH 8.0, 300 mM sucrose, 10 mM NaCl, 2 mM MgAc_2_ (all from Sigma). All following steps were performed at 4°C. Then, 500 μl of lysis buffer: 10 mM Tris-Cl pH 8.0, 300 mM sucrose, 10 mM NaCl, 2 mM MgAc_2_, 6 mM CaCl_2_, 0.2% NP-40/Igepal (all from Sigma), were added to the resuspended cells followed by pipetting the cells up and down 10 times. The total volume was then brought to 10 ml by adding 9 ml: 4.5 ml wash buffer, 4.5 ml lysis buffer. The tubes were mixed by inverting gently for 1 min, then nuclei were pelleted at 800xg for 5 min. The nuclei were then washed with 1ml of storage buffer: 50 mM Tris-HCL pH 8.3, 40% glycerol, 5 mM MgCl2, 0.1 mM EDTA (all from Sigma). Then, nuclei were counted and 5 x 10^6^ were pelleted per replicate in 1.5ml Eppendorf tubes. Pellets were resuspended in 42 μl storage buffer. A similar procedure was performed separately for *D.melanogaster* S2 cells. In the final step, 8 μl storage buffer with 35 x 10^3^ were added to the 42 μl storage buffer with mESC nuclei, and frozen in LN2 until the run-on reaction.

PRO-seq libraries were prepared according to (Mahat et al. 2016). Adjustments from the original protocol are: (1) In the Run-on Master Mix, the biotinylated nucleotides were provided at the following concentrations: 10 mM Biotin-11-ATP, 10 mM Biotin-11-GTP, 100 mM Biotin-11-CTP, and 100 mM Biotin-11-UTP; (2) Trizol LS (Life Technologies, 10296-010) was replaced by TRI Reagent-LS (MRC #TS 120); (3) Trizol (Life Technologies, 15596-026) was replaced by TRI Reagent (MRC #TR 118) ; (4) Digested RNA by base hydrolysis in 0.2 N NaOH on ice was reduced from 8_min to 6 min; (5) Nascent RNA was purified by binding streptavidin beads (NEB, S1421S) and washed as described. Hydrophilic Streptavidin Magnetic Beads (NEB, S1421S) was replaced by Streptavidin Magnetic Beads (NEB S1420S); (6) Superscript III Reverse Transcriptase (Life Technologies, 18080-044) was replaced by SuperScript IV Reverse Transcriptase (Life Tech. #18090050). Libraries were prepared using adapters that contain a 6bp unique molecular identifier sequence on read1.

### PRO-seq analysis

PRO-seq libraries were competitively aligned to a genome resulted by merging mm10 assembly with *D.melanogaster* dm3 genome assembly. Alignment was performed using the proseq2.0 pipeline developed by the Danko lab (https://github.com/Danko-Lab/proseq2.0) using the parameters -PE --RNA5=R2_5prime --UMI1=6. Downstream analysis was performed in R, using Genomic Ranges (Lawrence et al. 2013) and BRgenomics1.1.3 (https://mdeber.github.io/index.html).

To account for global changes in nascent RNA production, as well as technical variations between libraries, spike-in *D.melanogaster* S2 nuclei were used as internal control. The ration between fly:mouse nuclei was 1:150. As normalization, we divided the mouse reads in each sample by the total number of fly reads in the same sample.

We quantified changes in gene expression using the GENCODE v20 annotations in mouse. To compute differential expression between treatments we used DEseq2. First, we used un-normalized BigWigs to count the total number of reads around each TSS or within gene bodies of annotated GENCODE v20 genes. For TSSs, we took a 300bp window centered on gene start sites, while gene bodies were defined as the entirety of the gene excluding the first and last 300bp from TSS and TES respectively. Then, we provided the raw PRO-seq counts as input to DEseq2. We used the total number of *Drosophila* reads as scaling factors. For generating meta-profiles, *Drosophila* spike-in normalized counts were used. TSS meta-profiles in Figure 4B and 4C were aligned to mESCs START-seq data due to better accuracy than GENCODE v20 (Henriques et al. 2018).

### Heatmaps

Heatmaps were generated using *Drosophila* spike-in normalized reads. We sorted GENCODE v20 genes by length and depicted the number of spike-in normalized reads per 1kb bins from 1kb upstream the annotated TSS to 200kb downstream.

### dREG peaks

We called regulatory element peaks using dREG gateway developed by the Danko lab at the Baker Institute and Cornell University (Danko et al. 2015; Wang et al. 2019).

### Analysis of Micro-C data

Publicly available Micro-C data was downloaded from GSE130275 (Hsieh et al. 2020). HiC profiles were plotted using hicPlotTADs and used to define TAD boundaries for Nanog, Fgf5 (Wolff et al. 2020). Regulatory regions for Nanog and Fgf5 were called using the custom virtual4C script developed by the Danko lab (https://github.com/Danko-Lab/HS_transcription_regulation) using parameters -w 4000000 -b 5000 -q 30. The obtained regulatory regions were overlapped with dREG calls to define putative enhancers that are in contact with the four promoters of interest.

### Initiation-release rate estimation

To estimate initiation rates for each gene, method described in Siepel, 2021 was implemented (Siepel 2021). Specifically, initiation rate is estimated by

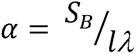

where *α* is initiation rate, *S_B_* is the number of read counts within gene body, *l* is gene length and *λ* is a library specific scaling factor determined by the number of spike-in reads mapped to *D. melanogaster* genome. While pause release rate is estimated by

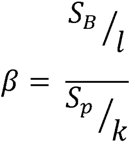

where is pause release rate, *S_p_* is the number of read counts within the pause peak and *k* is the length of it. The first protein coding annotations from GENCODE (version vM20) were used for each gene, and regions 1kb downstream of TSS to the end of the gene (up to 90kb) were used as gene body for read counting. Note *α* is the maximum likelihood estimator of initiation rate when assuming read counts following Poisson distribution, and it’s also widely used in many literatures to represent transcriptional activity with some heuristic justifications (Siepel 2021).

### Image processing and quantification

For Pre-implantation embryos, semi-automated 3D nuclear segmentation for cell counting and quantification of fluorescence intensity was carried out using MINS, a MATLAB-based algorithm (http://katlab-tools.org/) (Lou et al. 2014).The same imaging parameters were used for all experiments consisting of the same primary and secondary antibody combinations to minimize quantitative variance due to image acquisition. The MINS output was checked for over- or under-segmentation and tables were corrected manually using Image J (NIH, https://imagej.nih.gov/ij/). Under-segmented nuclei (two or more nuclei detected as one, or nuclei that were not detected) were assigned fluorescence intensity values that were directly measured using ImageJ (NIH). To correct fluorescence decay along the Z-axis, we used a linear regression method to calculate the global average of the regression coefficients in the HA channel(Saiz et al. 2016b). This slope was then used to adjust the logarithm values of HA fluorescence intensity for each nucleus. Trophectoderm (TE) vs. inner cell mass (ICM) cell assignment was achieved by a threshold for CDX2 which is present exclusively in TE. To avoid batch variability, directly compared embryos were stained and imaged in the same session.

### Statistical analysis

All statistical tests of immunofluorescence data were carried out in PRISM 9 (GraphPad). Statical significance was established using a student t-test with p-value threshold of 0.05. The p-value range for each experiment is indicated in the figure legend.

For sequencing data, analysis of differentially expressed genes was done in R using the DEseq2 method with 0.05 p. adjusted (Love et al. 2014).Other comparisons between gene groups were performed using two-way paired t-test.

### Data and code availability

Raw and processed sequencing data from this work is deposited in the Gene Expression Omnibus under the accession numbers GSE196543 for ChIP-seq and GSE196653 for PRO-seq.

## Reagents Table

| REAGENT | SOURCE | IDENTIFIER |
| --- | --- | --- |
| Antibodies |  |  |
| Antibodies in Table S1 | This paper | N/A |
| Bacterial and Virus Strains |  |  |
| E. coli DH5a | NEB | C2987I |
| Chemicals, Peptides, and Recombinant Proteins |  |  |
| Hoechst 33342 | Invitrogen | Cat# H3570 |
| Stemolecule™ CHIR99021 | Fisher | Cat# NC9785126 |
| PD0325901 | Reprocell | Cat# 04-0006-10 |
| dTAG-13 | Tocris Bio | Cat# 6605 |
| FHM | Millipore | Cat# MR-025-D |
| Proteinase K | Roche | Cat# 03115801001 |
| 16% PFA | Electron<br>microscopy sciences | Cat# 15710 |
| Acid Tyrode's | Millipore | Cat# MR-004-D |
| Glycine | Sigma | Cat# G7403 |
| Triton X-100 | Sigma | Cat# X100 |
| Horse Serum | Sigma | Cat# H0146 |
| DMEM | Life technologies | Cat# 11995073 |
| NEAA | Life technologies | Cat# 11140-050 |
| Glutamine | Life technologies | Cat# 25030164 |
| Sodium Pyruvate | Life technologies | Cat# 11360070 |
| 2-mercaptoethanol | Life technologies | Cat# 21985023 |
| Fetal Calf Serum | VWR | Cat# 97068-085 |
| 0.05% Trypsin EDTA | Life technologies | Cat# 25200114 |
| Gelatine | Millipore | Cat# 104070 |
| Penicillin/Streptomycin | Life technologies | Cat# 15140163 |
| Mitomycin C | Sigma | Cat# M4287 |
| Critical Commercial Assays |  |  |
| NEBuilder HiFi assembly kit | NEB | Cat# E5520S |
| Pierce<sup>TM</sup> BCA protein assay kit | Thermo | Cat# 23225 |
| Deposited Data |  |  |
| ChIP-seq | GEO | GSE196543 |
| PRO-seq | GEO | GSE196653 |
| Experimental Models: Cell Lines |  |  |
| Mouse: Embryonic stem cell<br>line E14 | ATCC | CRL-1821 |
| Experimental Models: Organisms/Strains |  |  |
| Mouse: CD1 | Charles River<br>Laboratory | Cat# 022 |
| Oligonucleotides |  |  |
| Oligonucleotides in Table S1 | This paper | N/A |
| Software and Algorithms |  |  |
| Fiji/ImageJ | (Schindelin et al.<br>2012) | https://imagej.nih.gov/ij/ |
| R Studio 1.1.456/R 4.0.2 | RStudio, Inc. | https://rstudio.com/ |
| DESeq2 3.13 | Bioconductor | https://bioconductor.org/packages/release/bioc/html/DESeq2.html |
| ggplot2 3.3.5 | R package | https://ggplot2.tidyverse.org |
| MINS | (Lou et al. 2014) | http://katlab-tools.org/ |
| Prism 9 | GraphPad<br>Software | https://www.graphpad.com/scientific-software/prism/ |
| ZEN | Carl Zeiss<br>Microsystems | https://www.zeiss.com/microscopy/us/products/microscope-software/zen.html |
| Other |  |  |
| Glass-bottom dish | MatTek | Cat# P35G-1.5-14-C |
| Nitrocellulose membrane | Cell Signaling | Cat #12369 |

## COMPETING INTEREST STATEMENT

The authors have no competing interests to declare.

## ACKNOWLEDGEMENTS

We thank MSK’s Integrated Genomics Operation and Bioinformatics Core Facility for assistance in sequencing and sequence data analysis, and Drs. Karen Adelman and Lucy Williams for sharing their *Nelfb^-/-^* mouse model. We are grateful to Dr. Effie Apostolou for providing feedback and guidance on this work, and members of the Hadjantonakis lab for stimulating discussion and critical feedback. AA is supported by a MSTP training grant from the NIH (T32GM007739) awarded to the Weill Cornell/Rockefeller/Sloan Kettering Tri-Institutional MD-PhD Program and NIH F30HD103398. Work is AS’s lab is supported by the NIH (R01HG010346 and R35GM127070). Work in CGD’s lab is supported by the NIH (R01HG009309) and NASA (17-EXO-17-2-0112). Work in AKH’s lab is supported by the NIH (R01HD094868, R01DK127821, R01HD086478, and P30CA008748).

## AUTHOR CONTRIBUTIONS

A.A and A.K.H. conceptualized the study. A.A. designed, performed, and analyzed most experiments. A.C. and C.G.D. analyzed and interpreted PRO-seq data. E.J.R. constructed PRO-seq libraries. Y.Z. and A.S. estimated the initiation-release rates. C.G.D supervised all bioinformatics analyses. A.A. wrote the manuscript with input from all authors. A.A., A.K.H, and C.G.D. acquired funding. A.K.H supervised the work.

## Notes

### Competing Interest Statement

The authors have declared no competing interest.

## REFERENCES

Abuhashem A, Garg V, Hadjantonakis A-K. 2022. RNA polymerase II pausing in development: orchestrating transcription. Open Biol 12: 210220.

Abuhashem A, Hadjantonakis A-K. 2021. Rapid and efficient adaptation of the dTAG system in mammalian development reveals stage specific requirements of NELF. *bioRxiv* 2021.11.30.470581. http://biorxiv.org/content/early/2021/11/30/2021.11.30.470581.abstract.

Adelman K, Lis JT. 2012. Promoter-proximal pausing of RNA polymerase II: emerging roles in metazoans. Nat Rev Genet 13: 720–731.

Amleh A, Nair SJ, Sun J, Sutherland A, Hasty P, Li R. 2009. Mouse cofactor of BRCA1 (Cobra1) is required for early embryogenesis. PLoS One 4: 2–9.

Aoi Y, Smith ER, Shah AP, Rendleman EJ, Marshall SA, Woodfin AR, Chen FX, Shiekhattar R, Shilatifard A. 2020. NELF Regulates a Promoter-Proximal Step Distinct from RNA Pol II Pause-Release. Mol Cell 78: 261–274.e5.

Chen FX, Smith ER, Shilatifard A. 2018. Born to run: control of transcription elongation by RNA polymerase II. Nat Rev Mol Cell Biol 19: 464–478.

Core L, Adelman K. 2019. Promoter-proximal pausing of RNA polymerase II: a nexus of gene regulation. Genes Dev 33: 960–982.

Core LJ, Waterfall JJ, Gilchrist DA, Fargo DC, Kwak H, Adelman K, Lis JT. 2012. Defining the status of RNA polymerase at promoters. Cell Rep 2: 1025–1035.

Cramer P. 2019. Organization and regulation of gene transcription. Nature 573: 45–54.

Danko CG, Hah N, Luo X, Martins AL, Core L, Lis JT, Siepel A, Kraus WL. 2013. Signaling pathways differentially affect RNA polymerase II initiation, pausing, and elongation rate in cells. Mol Cell 50: 212–222. https://pubmed.ncbi.nlm.nih.gov/23523369.

Danko CG, Hyland SL, Core LJ, Martins AL, Waters CT, Lee HW, Cheung VG, Kraus WL, Lis JT, Siepel A. 2015. Identification of active transcriptional regulatory elements from GRO-seq data. Nat Methods 12: 433–438.

Gilchrist DA, Dos Santos G, Fargo DC, Xie B, Gao Y, Li L, Adelman K. 2010. Pausing of RNA polymerase II disrupts DNA-specified nucleosome organization to enable precise gene regulation. Cell 143: 540–551.

Gilchrist DA, Fromm G, dos Santos G, Pham LN, McDaniel IE, Burkholder A, Fargo DC, Adelman K. 2012. Regulating the regulators: the pervasive effects of Pol II pausing on stimulus-responsive gene networks. Genes Dev 26: 933–944. https://pubmed.ncbi.nlm.nih.gov/22549956.

Gressel S, Schwalb B, Cramer P. 2019. The pause-initiation limit restricts transcription activation in human cells. Nat Commun 10: 3603.

Hah N, Murakami S, Nagari A, Danko CG, Kraus WL. 2013. Enhancer transcripts mark active estrogen receptor binding sites. Genome Res 23: 1210–1223.

Hamilton WB, Mosesson Y, Monteiro RS, Emdal KB, Knudsen TE, Francavilla C, Barkai N, Olsen J V, Brickman JM. 2019. Dynamic lineage priming is driven via direct enhancer regulation by ERK. Nature 575: 355–360. https://doi.org/10.1038/s41586-019-1732-z.

Hayashi K, Ohta H, Kurimoto K, Aramaki S, Saitou M. 2011. Reconstitution of the mouse germ cell specification pathway in culture by pluripotent stem cells. Cell 146: 519–532.

Henriques T, Gilchrist DA, Nechaev S, Bern M, Muse GW, Burkholder A, Fargo DC, Adelman K. 2013. Stable pausing by RNA polymerase II provides an opportunity to target and integrate regulatory signals. Mol Cell 52: 517–528. https://pubmed.ncbi.nlm.nih.gov/24184211.

Henriques T, Scruggs BS, Inouye MO, Muse GW, Williams LH, Burkholder AB, Lavender CA, Fargo DC, Adelman K. 2018. Widespread transcriptional pausing and elongation control at enhancers. Genes Dev 32: 26–41.

Hewitt SC, Li R, Adams N, Winuthayanon W, Hamilton KJ, Donoghue LJ, Lierz SL, Garcia M, Lydon JP, DeMayo FJ, et al. 2019. Negative elongation factor is essential for endometrial function. FASEB J Off Publ Fed Am Soc Exp Biol 33: 3010–3023.

Hsieh T-HS, Cattoglio C, Slobodyanyuk E, Hansen AS, Rando OJ, Tjian R, Darzacq X. 2020. Resolving the 3D Landscape of Transcription-Linked Mammalian Chromatin Folding. Mol Cell 78: 539–553.e8.

Johnston RJJ, Desplan C. 2010. Stochastic mechanisms of cell fate specification that yield random or robust outcomes. Annu Rev Cell Dev Biol 26: 689–719.

Kim T-K, Hemberg M, Gray JM, Costa AM, Bear DM, Wu J, Harmin DA, Laptewicz M, Barbara-Haley K, Kuersten S, et al. 2010. Widespread transcription at neuronal activity-regulated enhancers. Nature 465: 182–187.

Krebs AR, Imanci D, Hoerner L, Gaidatzis D, Burger L, Schübeler D. 2017. Genome-wide Single-Molecule Footprinting Reveals High RNA Polymerase II Turnover at Paused Promoters. Mol Cell 67: 411–422.e4.

Kwak H, Fuda NJ, Core LJ, Lis JT. 2013. Precise maps of RNA polymerase reveal how promoters direct initiation and pausing. Science 339: 950–953. https://pubmed.ncbi.nlm.nih.gov/23430654.

Kwon GS, Fraser ST, Eakin GS, Mangano M, Isern J, Sahr KE, Hadjantonakis A-K, Baron MH. 2006. Tg(Afp-GFP) expression marks primitive and definitive endoderm lineages during mouse development. Dev Dyn an Off Publ Am Assoc Anat 235: 2549–2558.

Lawrence M, Huber W, Pagès H, Aboyoun P, Carlson M, Gentleman R, Morgan MT, Carey VJ. 2013. Software for computing and annotating genomic ranges. PLoS Comput Biol 9: e1003118.

Liu X, Kraus WL, Bai X. 2015. Ready, pause, go: regulation of RNA polymerase II pausing and release by cellular signaling pathways. Trends Biochem Sci 40: 516–525.

Lou X, Kang M, Xenopoulos P, Munoz-Descalzo S, Hadjantonakis A-K. 2014. A rapid and efficient 2D/3D nuclear segmentation method for analysis of early mouse embryo and stem cell image data. Stem cell reports 2: 382–397.

Love MI, Huber W, Anders S. 2014. Moderated estimation of fold change and dispersion for RNA-seq data with DESeq2. Genome Biol 15: 550. https://doi.org/10.1186/s13059-014-0550-8.

Mahat DB, Kwak H, Booth GT, Jonkers IH, Danko CG, Patel RK, Waters CT, Munson K, Core LJ, Lis JT. 2016. Base-pair-resolution genome-wide mapping of active RNA polymerases using precision nuclear run-on (PRO-seq). Nat Protoc 11: 1455. https://doi.org/10.1038/nprot.2016.086.

Morgani S, Nichols J, Hadjantonakis A-K. 2017. The many faces of Pluripotency: in vitro adaptations of a continuum of in vivo states. BMC Dev Biol 17: 7. https://doi.org/10.1186/s12861-017-0150-4.

Nabet B, Roberts JM, Buckley DL, Paulk J, Dastjerdi S, Yang A, Leggett AL, Erb MA, Lawlor MA, Souza A, et al. 2018. The dTAG system for immediate and target-specific protein degradation. Nat Chem Biol 14: 431–441. https://doi.org/10.1038/s41589-018-0021-8.

Narita T, Yung TMC, Yamamoto J, Tsuboi Y, Tanabe H, Tanaka K, Yamaguchi Y, Handa H. 2007. NELF interacts with CBC and participates in 3’ end processing of replication-dependent histone mRNAs. Mol Cell 26: 349–365.

Nechaev S, Fargo DC, dos Santos G, Liu L, Gao Y, Adelman K. 2010. Global analysis of short RNAs reveals widespread promoter-proximal stalling and arrest of Pol II in Drosophila. Science 327: 335–338.

Ou J, Guan X, Wang J, Wang T, Zhang B, Li R, Xu H, Hu X, Guo X-K. 2021. Epithelial NELF guards intestinal barrier function to ameliorate colitis by maintaining junctional integrity. Mucosal Immunol.

Pope SD, Medzhitov R. 2018. Emerging Principles of Gene Expression Programs and Their Regulation. Mol Cell 71: 389–397.

Ran FA, Hsu PD, Wright J, Agarwala V, Scott DA, Zhang F. 2013. Genome engineering using the CRISPR-Cas9 system. Nat Protoc 8: 2281–2308.

Robinson DCL, Ritso M, Nelson GM, Mokhtari Z, Nakka K, Bandukwala H, Goldman SR, Park PJ, Mounier R, Chazaud B, et al. 2021. Negative elongation factor regulates muscle progenitor expansion for efficient myofiber repair and stem cell pool repopulation. Dev Cell 56: 1014–1029.e7.

Rougvie AE, Lis JT. 1988. The RNA polymerase II molecule at the 5’ end of the uninduced hsp70 gene of D. melanogaster is transcriptionally engaged. Cell 54: 795–804.

Saiz N, Kang M, Schrode N, Lou X, Hadjantonakis A-K. 2016a. Quantitative Analysis of Protein Expression to Study Lineage Specification in Mouse Preimplantation Embryos. J Vis Exp 53654.

Saiz N, Williams KM, Seshan VE, Hadjantonakis A-K. 2016b. Asynchronous fate decisions by single cells collectively ensure consistent lineage composition in the mouse blastocyst. Nat Commun 7: 13463. https://doi.org/10.1038/ncomms13463.

Schindelin J, Arganda-Carreras I, Frise E, Kaynig V, Longair M, Pietzsch T, Preibisch S, Rueden C, Saalfeld S, Schmid B, et al. 2012. Fiji: an open-source platform for biological-image analysis. Nat Methods 9: 676–682.

Schwalb B, Michel M, Zacher B, Frühauf K, Demel C, Tresch A, Gagneur J, Cramer P. 2016. TT-seq maps the human transient transcriptome. Science 352: 1225–1228.

Shao R, Kumar B, Lidschreiber K, Lidschreiber M, Cramer P, Elsässer SJ. 2021. Newly synthesized RNA Sequencing Characterizes Transcription Dynamics in Three Pluripotent States. bioRxiv 2021.06.11.448016. http://biorxiv.org/content/early/2021/06/13/2021.06.11.448016.abstract.

Shao W, Zeitlinger J. 2017. Paused RNA polymerase II inhibits new transcriptional initiation. Nat Genet 49: 1045–1051. https://doi.org/10.1038/ng.3867.

Siepel A. 2021. A Unified Probabilistic Modeling Framework for Eukaryotic Transcription Based on Nascent RNA Sequencing Data. *bioRxiv* 2021.01.12.426408. http://biorxiv.org/content/early/2021/01/14/2021.01.12.426408.abstract.

Steurer B, Janssens RC, Geverts B, Geijer ME, Wienholz F, Theil AF, Chang J, Dealy S, Pothof J, van Cappellen WA, et al. 2018. Live-cell analysis of endogenous GFP-RPB1 uncovers rapid turnover of initiating and promoter-paused RNA Polymerase II. Proc Natl Acad Sci U S A 115: E4368–E4376.

Vos SM, Farnung L, Boehning M, Wigge C, Linden A, Urlaub H, Cramer P. 2018a. Structure of activated transcription complex Pol II-DSIF-PAF-SPT6. Nature 560: 607–612.

Vos SM, Farnung L, Urlaub H, Cramer P. 2018b. Structure of paused transcription complex Pol II–DSIF–NELF. *Nature*. http://www.nature.com/articles/s41586-018-0442-2.

Wang X, Hang S, Prazak L, Gergen JP. 2010. NELF Potentiates Gene Transcription in the Drosophila Embryo. PLoS One 5: e11498. https://doi.org/10.1371/journal.pone.0011498.

Wang Z, Chu T, Choate LA, Danko CG. 2019. Identification of regulatory elements from nascent transcription using dREG. Genome Res 29: 293–303.

Whyte WA, Orlando DA, Hnisz D, Abraham BJ, Lin CY, Kagey MH, Rahl PB, Lee TI, Young RA. 2013. Master transcription factors and mediator establish super-enhancers at key cell identity genes. Cell 153: 307–319. https://www.ncbi.nlm.nih.gov/pubmed/23582322.

Williams LH, Fromm G, Gokey NG, Henriques T, Muse GW, Burkholder A, Fargo DC, Hu G, Adelman K. 2015. Pausing of RNA Polymerase II Regulates Mammalian Developmental Potential through Control of Signaling Networks. Mol Cell 58: 311–322. http://dx.doi.org/10.1016/j.molcel.2015.02.003.

Wissink EM, Vihervaara A, Tippens ND, Lis JT. 2019. Nascent RNA analyses: tracking transcription and its regulation. Nat Rev Genet 20: 705–723. https://pubmed.ncbi.nlm.nih.gov/31399713.

Wolff J, Rabbani L, Gilsbach R, Richard G, Manke T, Backofen R, Grüning BA. 2020. Galaxy HiCExplorer 3: a web server for reproducible Hi-C, capture Hi-C and single-cell Hi-C data analysis, quality control and visualization. Nucleic Acids Res 48: W177–W184. https://pubmed.ncbi.nlm.nih.gov/32301980.

Wu C-H, Yamaguchi Y, Benjamin LR, Horvat-Gordon M, Washinsky J, Enerly E, Larsson J, Lambertsson A, Handa H, Gilmour D. 2003. NELF and DSIF cause promoter proximal pausing on the hsp70 promoter in Drosophila. Genes Dev 17: 1402–1414.

Wu T, Hadjantonakis A-K, Nowotschin S. 2017. Visualizing endoderm cell populations and their dynamics in the mouse embryo with a Hex-tdTomato reporter. Biol Open 6: 678–687.

Wu T, Yoon H, Xiong Y, Dixon-Clarke SE, Nowak RP, Fischer ES. 2020. Targeted protein degradation as a powerful research tool in basic biology and drug target discovery. Nat Struct Mol Biol 27: 605–614.

Yamaguchi Y, Takagi T, Wada T, Yano K, Furuya A, Sugimoto S, Hasegawa J, Handa H. 1999. NELF, a multisubunit complex containing RD, cooperates with DSIF to repress RNA polymerase II elongation. Cell 97: 41–51.

Yang Q, Liu X, Zhou T, Cook J, Nguyen K, Bai X. 2016. RNA polymerase II pausing modulates hematopoietic stem cell emergence in zebrafish. Blood 128: 1701–1710. https://pubmed.ncbi.nlm.nih.gov/27520065.

Yu L, Zhang B, Deochand D, Sacta MA, Coppo M, Shang Y, Guo Z, Zeng X, Rollins DA, Tharmalingam B, et al. 2020. Negative elongation factor complex enables macrophage inflammatory responses by controlling anti-inflammatory gene expression. Nat Commun 11: 2286. https://doi.org/10.1038/s41467-020-16209-5.

