## Supplementary material for "RNA Pol II pausing facilitates phased pluripotency transitions by buffering transcription": Figure S1

Abuhashem_Fig S1

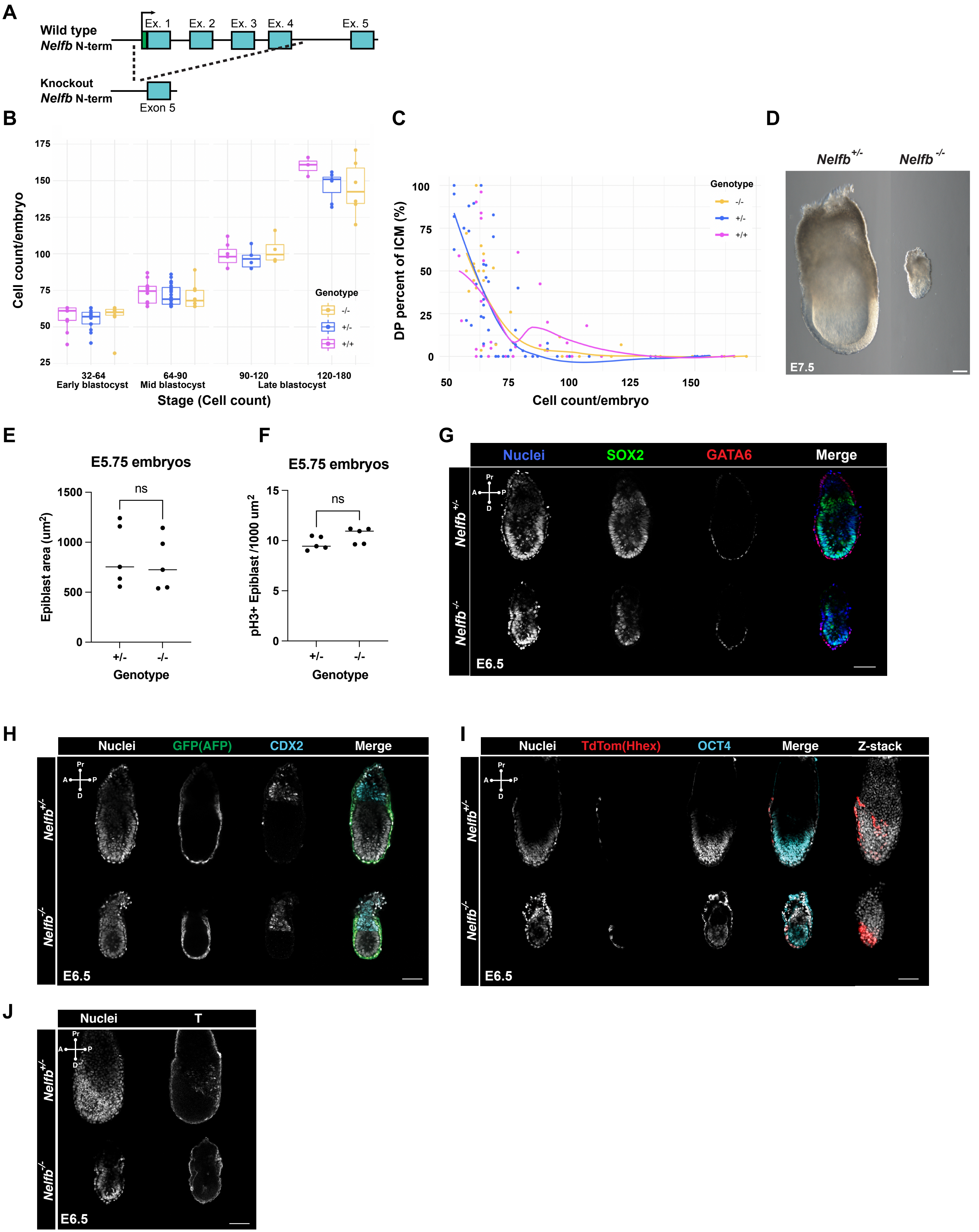

**Fig S1.** ***Nelfb*$^{-/-}$ embryos display defects in pluripotent epiblast state transitions, related to Figure 1.**

(A) Schematic of the *Nelfb* knockout (*Nelfb*$^{-}$) allele.

(B) Total cell count per blastocyst per genotype per stage.

(C) Percentage of double positive nuclei: NANOG+ GATA6+ of the ICM in each blastocyst per genotype. Each dot represents an embryo. Loess line fit is shown.

(D) Brightfield images of heterozygous and null embryos at E7.5. Scale bar 100µm

(E) Epiblast layer area in E5.75 embryos.

(F) Ratio (count/area) of dividing nuclei in the epiblast cup region. Dividing nuclei were visualized via immunostaining for phosphorylated H3 (pH3). Statistical testing was performed using a t-test.

(G – J) Immunofluorescence of E6.5 embryos. Nuclei are labeled with Hoechst. Single Z slices are shown. Scale bar 100µm.
