## Supplementary material for "RNA Pol II pausing facilitates phased pluripotency transitions by buffering transcription": Figure S2

### Abuhashem_Fig S2

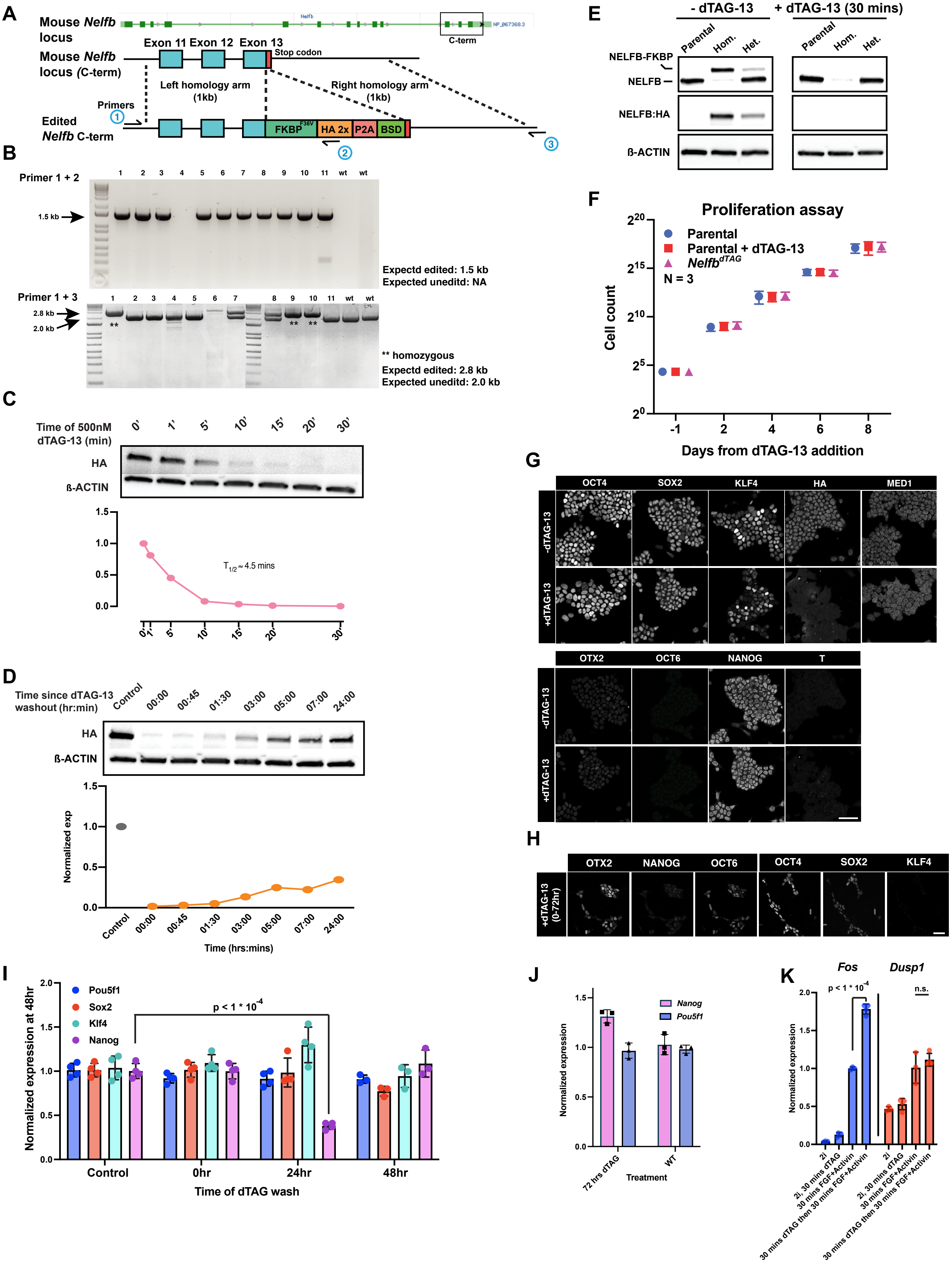

**Fig S2. NELFB-depleted mESCs recapitulate defects in pluripotent state transitions observed in the embryo, related to Figure 2.**

(A) Top: NCBI genome browser shot of mouse *Nelfb* locus. Bottom: Schematic of *Nelfb*$^{deg}$ knock-in strategy.

(B) Genotyping of targeted clones to identify correctly targeted homozygous clones.

(C) Western blot and quantification of NELFB degradation dynamics with 5 mins intervals.

(D) Western blot and quantification of NELFB recovery dynamics following 1 hour dTAG-13 treatment and wash.

(E) Western blot using NELFB antibody to validate correctly generated homozygous clones.

(F) Proliferation assay of parental cells and *Nelfb*$^{deg}$ mESC in the presence and absence of dTAG-13. Cells were counted and passaged every two days. Control conditions Related to Figure 2D.

(G) Immunofluorescence of *Nelfb*$^{deg}$ mESC at 0hr of transitions representing naïve state with and without 24 hours of dTAG-13. Scale bar 50µm.

(H) Immunofluorescence of *Nelfb*$^{deg}$ mESC following pluripotency transitions at 72 hours with dTAG-13 from 0-72 hours. Related to Figure 2F. Scale bar 50µm.

(I) Normalized RT-qPCR expression of select genes with transient dTAG-13 treatment, 1 hour followed by wash. Data points show only hour 48 analysis for statistical validation. Related to Figure 2H. Statistical testing was performed using a t-test.

(J) Normalized RT-qPCR expression of select genes with dTAG-13 treatment in naïve *Nelfb*$^{deg}$ mESC (0hr state) for 72 hours. Statistical testing was performed using a t-test.

(K) Normalized RT-qPCR expression of select genes with 30 mins dTAG-13 treatment in naïve *Nelfb*$^{deg}$ mESC (0hr state) followed by acute 30 mins FGF + ACTIVIN treatment. Statistical testing was performed using a t-test.

All RT-qPCR data is normalized to *Actb*, then control.
