## Supplementary material for "RNA Pol II pausing facilitates phased pluripotency transitions by buffering transcription": Figure S3

### Abuhashem_Fig S3

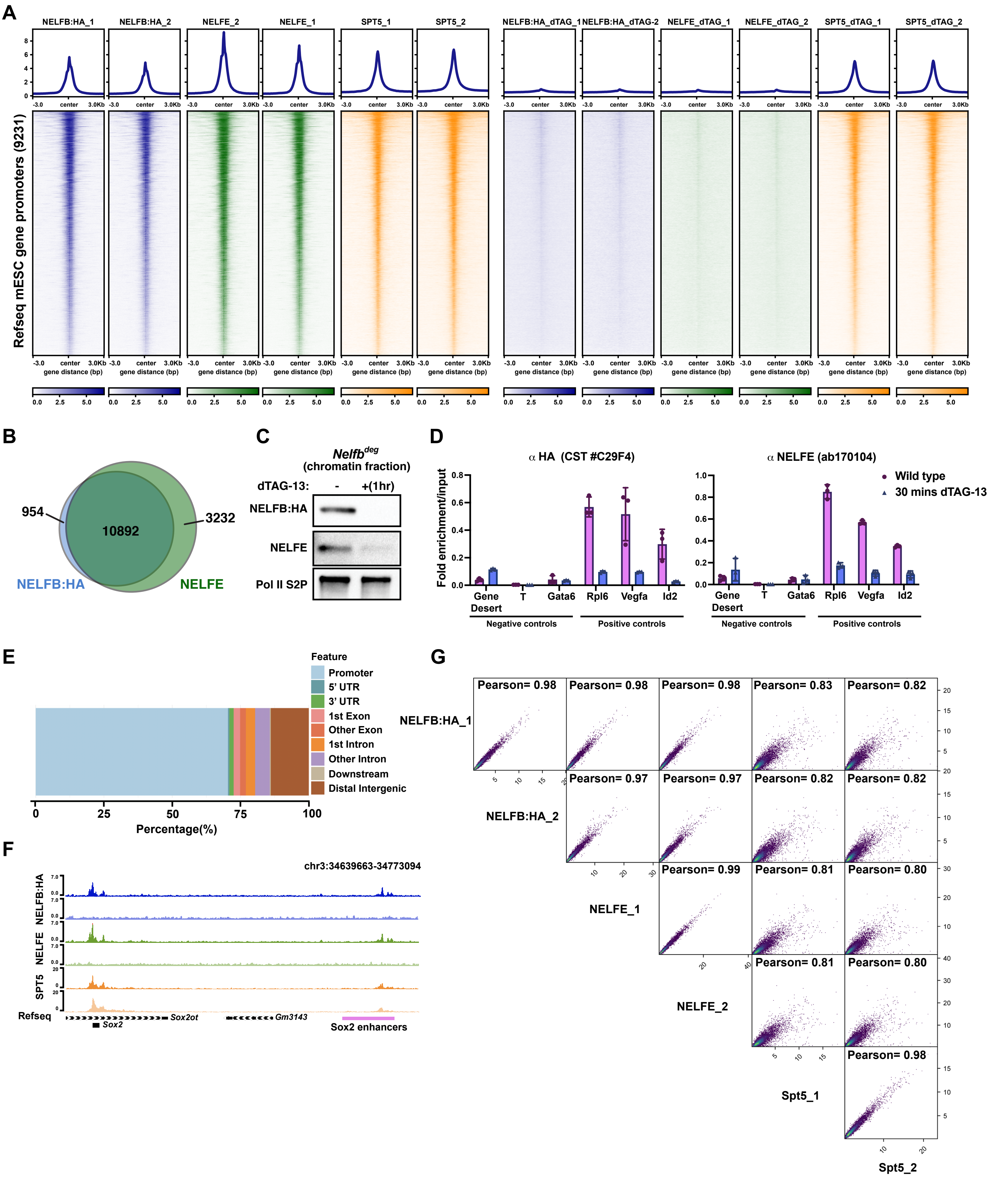

**Fig S3. NELF displays widespread binding at promoters and enhancers and *Nelfb*$^{deg}$ enables acute clearance of the NELF complex from chromatin.**

(A) Heatmaps of NELFB, NELFE, and SPT5 signal across replicates and conditions presented in Figure 3A.

(B) Venn diagram of overlap between NELFB and NELFE peaks. Peaks have a q. value < 0.05 cutoff.

(C) Western blot of chromatin fraction *Nelfb*$^{deg}$ mESC with and without 1 hour of dTAG-13 treatment.

(D) ChIP-qPCR validation of ChIP experiments NELFB and NELFE following 30 mins of dTAG-13 treatment.

(E) NELF peaks annotation.

(F) Genome browser shot of a representative enhancer region showing NELF peaks. Related to Figure 3F.

(G) Signal Pearson correlation across ChIP-seq replicates and samples.
