## Supplementary material for "RNA Pol II pausing facilitates phased pluripotency transitions by buffering transcription": Figure S4

### Abuhashem_Fig S4

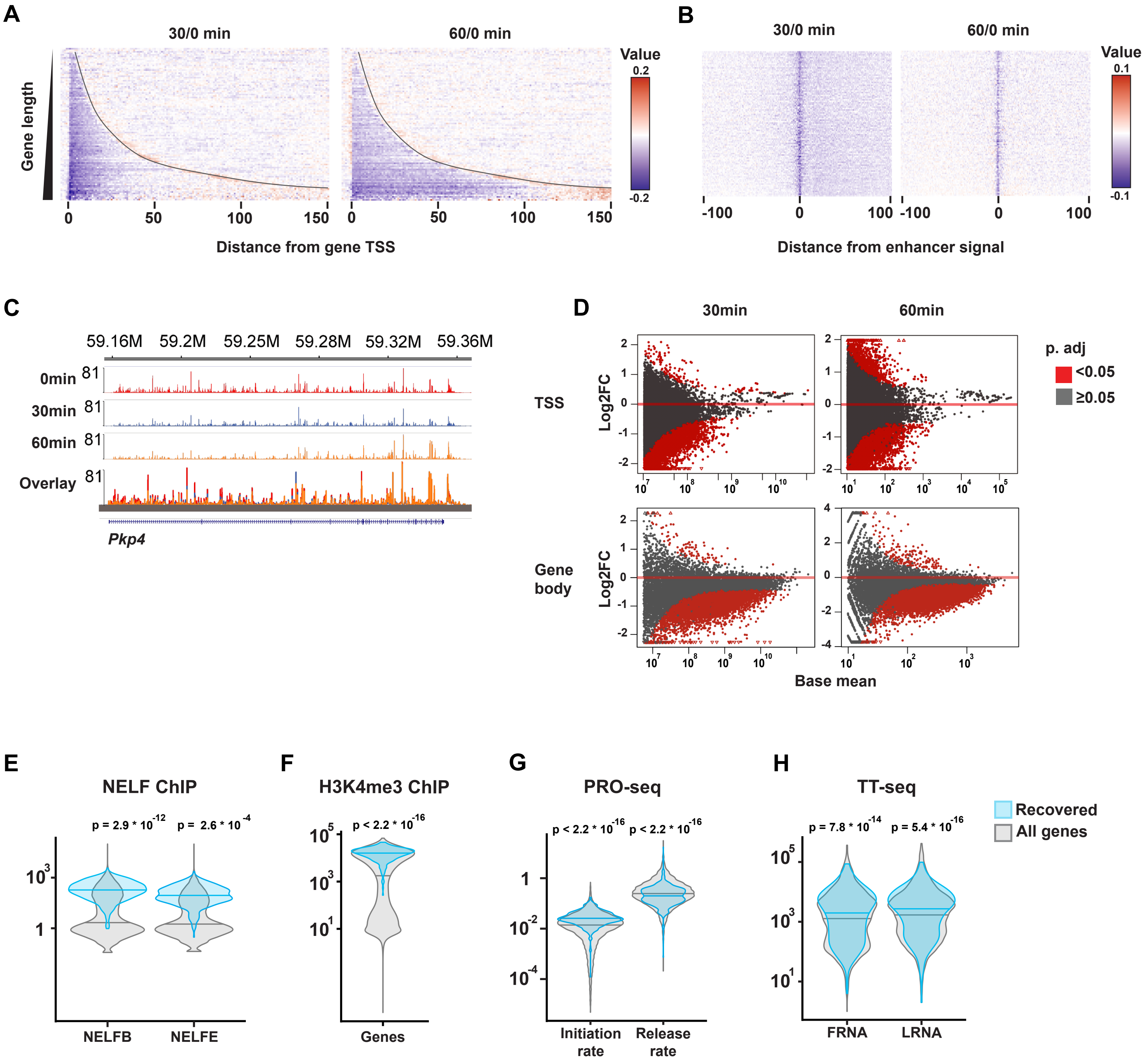

**Fig S4. NELF stabilizes Pol II pausing and transcription at promoters and enhancers, related to Figure 4.**

(A) Heatmaps of Log2 fold change of PRO-seq signal at genes ordered by gene length. Dashed lines represent poly A sites.

(B) Heatmaps of Log2 fold change of PRO-seq signal at enhancers.

(C) Genome browser shot of PRO-seq signal for a long gene, related to Figure 4E.

(D) MA plot of Log2 fold change with significantly changed genes highlighted. Related to Figure 4F and 4H.

(E) Transcription initiation and release rates from PRO-seq data at 0 mins of all genes vs. recovering TSS genes at 60 mins treatment.

(F) NELF ChIP signal at all genes vs. recovering TSS genes at 60 mins treatment.

(G) H3K4me3 signal at all genes vs. recovering TSS genes at 60 mins treatment.

(H) TT-seq signal at all genes vs. recovering TSS genes at 60 mins treatment.
