## Supplementary material for "RNA Pol II pausing facilitates phased pluripotency transitions by buffering transcription": Figure S5

### Abuhashem_Fig S5

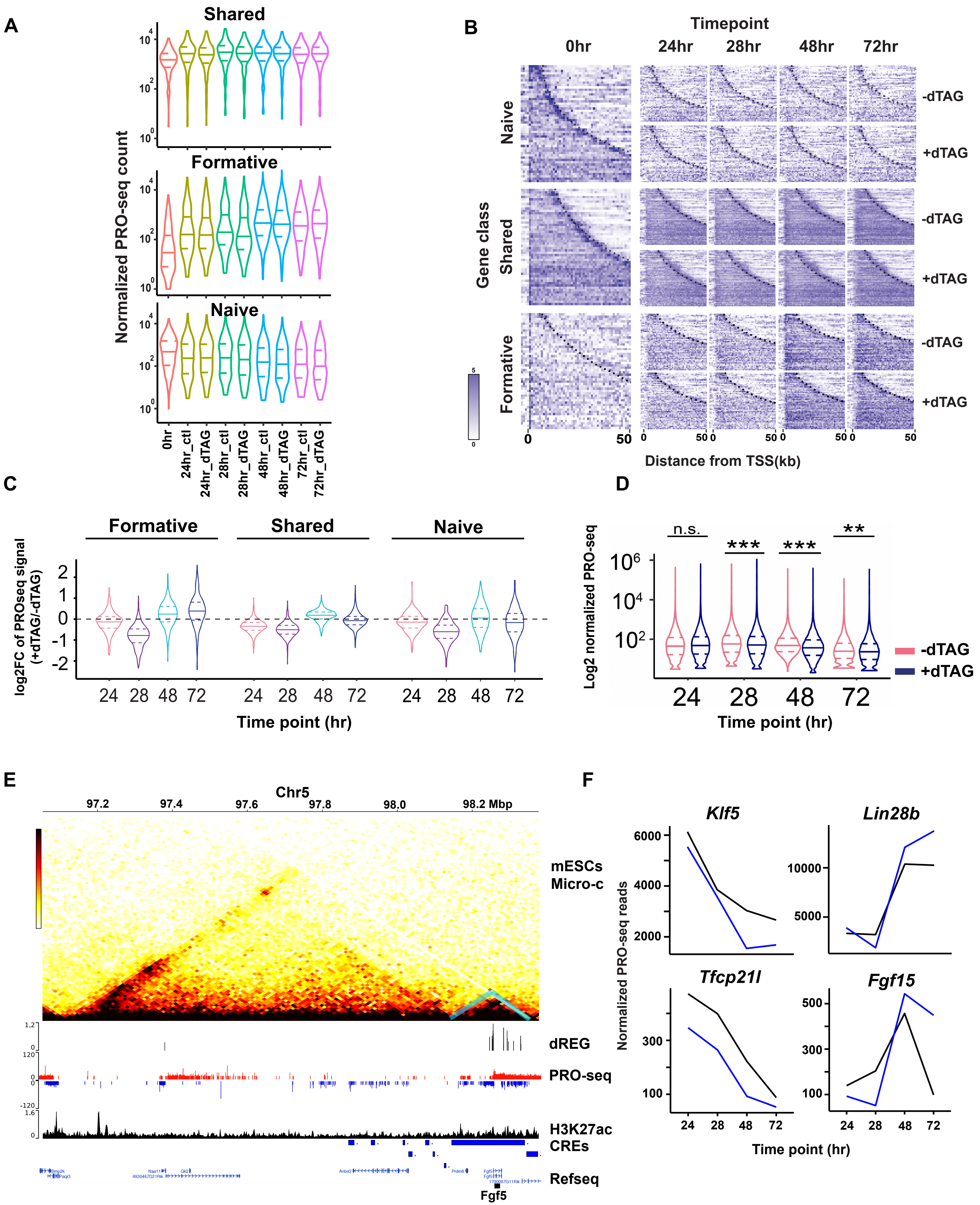

**Fig S5. NELF balances gene induction and repression during pluripotency transitions, related to Figure 5**

(A) Violin plot of normalized PRO-seq count/gene in each gene class per treatment per time point in our analysis, related to Figure 5C.

(B) Heatmap of PRO-seq signal in each gene class at each time point with and without dTAG-13. Dashed lines represent poly A sites, related to Figure 5C and S5A.

(C) Violin plot of log2 fold change of PRO-seq data gene expression at each time point of the analysis per gene group, related to Figure 5E.

(D) Violin plot of PRO-seq signal at TREs identified by dREG at each time point with and without dTAG-13.

(E) Example of defining putative enhancers for certain loci using public Micro-c data, dREG, and H3K27ac marks. Fgf5 locus is shown. Related to Figure 5F.

(F) PRO-seq signal of additional naïve genes (Klf5 and Tfcp21l) and formative genes (Lin28b and Fgf15) during transitions, with and without dTAG-13 treatments, related to Figure 5F.
