## Supplementary material for "RNA Pol II pausing facilitates phased pluripotency transitions by buffering transcription": Figure S6

### Abuhashem_Fig S6

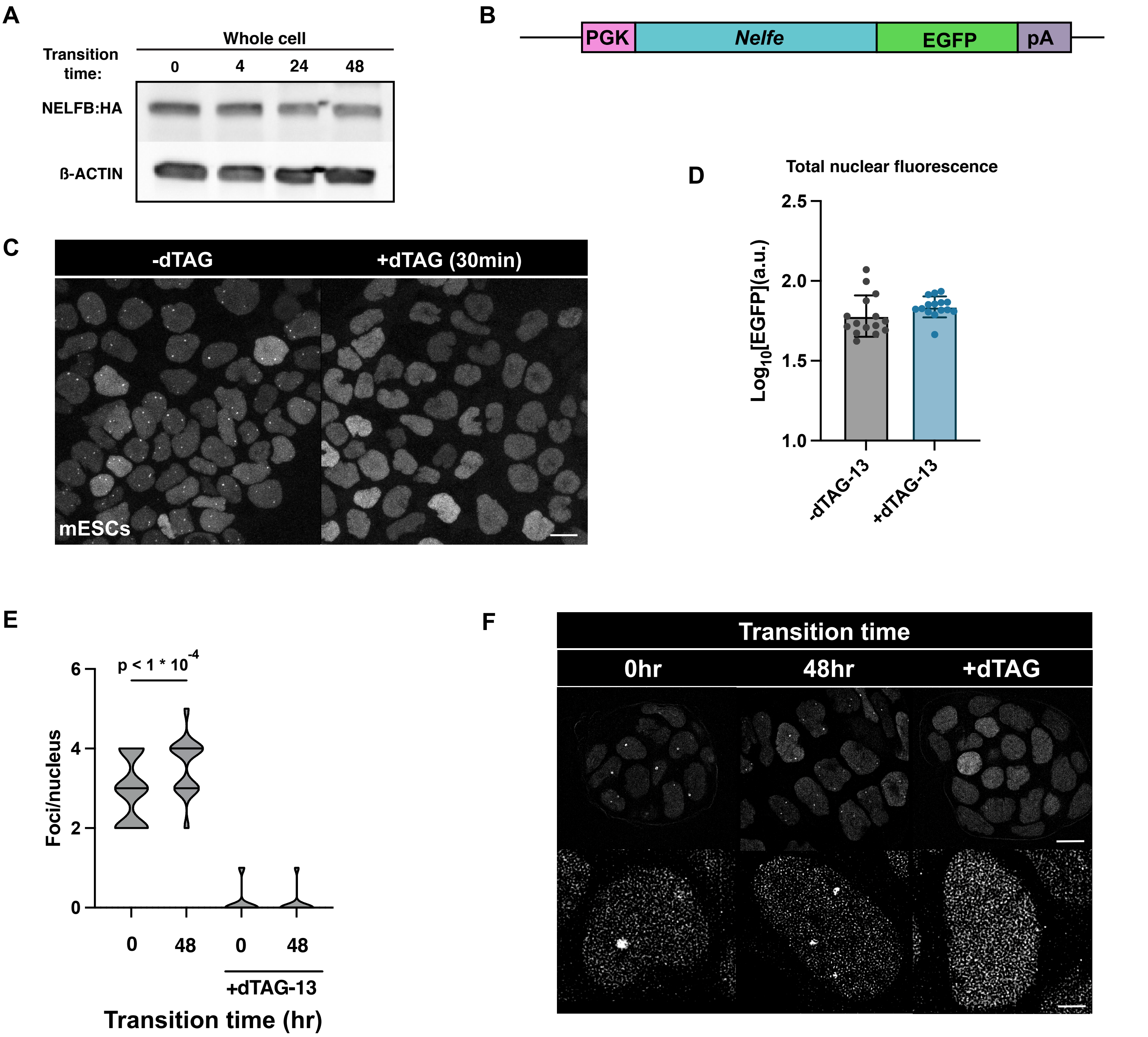

**Fig S6. NELF is recruited to chromatin during pluripotency transitions, related to Figure 6**

(A) Western blot of whole cell lysates during pluripotency transitions, related to Figure 6A.

(B) Nelfe-EGFP transgene construct design.

(C) Live images of NELFE-EGFP with and without 30 mins of dTAG-13. Scale bar 10μm.

(D) Whole nucleus EGFP fluorescence quantification of images in Figure S6C.

(E) Violin plot of NELF bodies per nuclei using super-resolution microscopy.

(F) Live super-resolution images of NELFE-EGFP. Scale bar: top 10μm; bottom 2μm.
